# Autonomous Reef Monitoring Structures (ARMS) Reveal Human-Induced Biodiversity Shifts in Urban Coastal Ecosystems

**DOI:** 10.1101/2025.07.11.664282

**Authors:** Zhongyue Wan, Isis Guibert, Wing Yi Haze Chung, Charlotte Ho, Wan Ching Rachel Au, Joseph Brennan, Ling Fung Matt Chan, Emily Chei, Inga Conti-Jerpe, Alison Corley, Jonathan D. Cybulski, Róisín Hayden, Shan Yee Joyce Lee, Wendy McLeod, Philip Thompson, Shelby E. McIlroy, David M. Baker

**Affiliations:** The Swire Institute of Marine Science, The University of Hong Kong, Cape D’Aguilar Road, Shek O, Hong Kong SAR; School of Biological Sciences, The University of Hong Kong, Pok Fu Lam, Hong Kong SAR; Simon F. S. Li Marine Science Laboratory, School of Life Sciences, The Chinese University of Hong Kong, Shatin, Hong Kong SAR; Department of Biology, University of Oxford, Oxford, UK; Tree of Life, Wellcome Sanger Institute, Hinxton, Cambridge, UK; Lingnan University, Tuen Mun, Hong Kong SAR

## Abstract

Biodiversity thrives in coastal marine habitats which host foundational species such as corals, mangroves, and seagrasses. However, coastal development and the growth of megacities along shorelines impose an array of stressors on the marine environment. These stressors inevitably impact biodiversity which dictates ecosystem functions and services. Despite extensive research on biodiversity responses to anthropogenic stressors, phylum-specific resistance and resilience dynamics – particularly in coastal marine ecosystems – remain poorly understood. Considering the global scale of coastal development, it is imperative to develop a more comprehensive understanding of how biodiversity, in terms of richness and community composition, is influenced by various anthropogenic stressors. Here, we present the first application of standardized Autonomous Reef Monitoring Structures (ARMS) as an experimental unit - using a common garden experimental design - to examine community responses to stress within an urbanized seascape. ARMS were seeded within two marine reserves for one year and then transplanted to sites of stress, including domestic sewage, and mariculture. We hypothesized that 1) human impacts reduce richness and alter composition of established communities; and 2) increasing intensity of these impacts reduces community resistance and resilience to stress. Using metabarcoding, we quantified richness and taxonomic composition and assessed their changes along an impact gradient. Our results showed that nutrient pollution, particularly inorganic nitrogen, significantly reduced species richness and restructured communities. Communities exhibited low resistance, yet high resilience - suggesting that urbanized seascapes have high recovery potential when stress is mitigated.

## 1. Introduction

It is estimated that more than 8 million species coexist on Earth, each playing a unique role in the intricate web of life (Wiens, 2023). This rich biodiversity underpins ecosystem functions that drive the flow of energy and the cycling of matter across space and time (Snelgrove et al., 2014). For example, marine algae convert solar energy into biomass via photosynthesis, fueling food webs; sponges enhance nutrient cycling in aquatic ecosystems through filter feeding; and detritivores decompose organic matter, returning essential elements to the seafloor and sustaining nutrient availability. Many ecosystem functions, through their interactions, directly benefit humans with economic value and are collectively termed “ecosystem services” (Mace et al., 2012). In marine ecosystems, these services are particularly vital: global annual seafood harvests average 50 million metric tons (MT), contributing to about 17% of the global meat production (Free et al., 2022); intricate microbial communities in marine sediments denitrify the ocean, returning over 200 teragrams (Tg) of nitrogen back to the atmosphere, alleviating eutrophication (Devol, 2015); the ocean absorbs about 25% of human-generated CO_2_, and marine organisms collectively produce over 50% of the O_2_ we breathe (Devries, 2024; Hocke, 2023).

Despite the importance of ecosystem functions and services, biodiversity faces unprecedented anthropogenic threats. Human activities drive planetary-scale environmental change, placing Earth on a trajectory toward its sixth mass extinction event (Cowie et al., 2022). Across marine ecosystems, the severity of extinction is intensifying due to the combined pressures of overfishing, nutrient pollution, and human-induced climate change (O’hara et al., 2021). This loss of biodiversity has diminished services across ecosystems. For example, following the decline of coral-reef-associated biodiversity, the coral reef fishery catch-per-unit-effort (total catch normalized by catching effort) has decreased since 1971 (Eddy et al., 2021), the global destruction of mangroves between 2000 and 2015 led to a release of over 30 Tg of soil carbon (Sanderman et al., 2018), and regional marine biodiversity loss has impaired detoxification and filtration services, leading to effects such as water quality degradation, harmful algal blooms (HAB), mass fish mortality events, and beach closures (Worm et al., 2006). These multilayer, cascading impacts underscore the urgency to study how human activities are reshaping biodiversity and destabilizing ecosystems.

The complex nexus between biodiversity, disturbance, and ecosystem stability is a cornerstone of ecology. For over 70 years, theoretical and empirical work has provided competing perspectives on biodiversity’s role in ecosystem stability under disturbance (Donohue et al., 2016; Van Meerbeek et al., 2021). Early ecologists predicted a positive relationship between species diversity and ecosystem stability. MacArthur (1955) proposed that a higher diversity of both prey and predator species enriched the complexity of the food-web with a stabilizing effect on the ecosystem; Elton (1958) further suggested that diversity can reduce the impact of invasive species and population explosions as a mechanism for increasing stability. These early ideas were challenged by May (1974) who created mathematical models with mock communities and found that higher richness tended to destabilize ecosystems – “*complexity begets instability*”. However, Yodzis (1981) questioned this viewpoint, discovering that modelled communities based on real food webs exhibited greater stability than randomized simulations. This debate has continued to this day, with recent studies showing mixed diversity-stability relationships that can be positive (Loreau & de Mazancourt, 2013), negative (Pennekamp et al., 2018), or exert opposing impacts on different levels of a system (Xu et al., 2021).

This field of study is further complicated by the increasingly tangled lexicon surrounding stability. The term “stability” and its several major variables have been repeatedly re-introduced and re-defined ambiguously by various stakeholders. Grimm and Wissel (1996) conducted a meta-analysis and found that 70 stability concepts were outlined with a total of 163 definitions. Terminological inconsistency remains a challenge. Stability has been assessed using both functional and compositional approaches (Donohue et al., 2013; Van Meerbeek et al., 2021). A system’s ability to resist disturbance is inconsistently labeled—either as *robustness* (Donohue et al., 2016) or *tolerance* (Van Meerbeek et al., 2021). Similarly, definitions of *resilience* vary: some emphasize recovery to a reference state (Grimm & Wissel, 1996), while others focus on the speed of recovery (Van Meerbeek et al., 2021). These discrepancies emphasize the need for conceptual clarity while investigating the diversity-stability relationship. In the following work, we employed the framework from Grimm & Wissel (1996) and centered on two main variables: 1) resistance: a community’s ability to resist changes during a period of environmental disturbance, and 2) resilience: the extent to which a community returns to the reference state after a disturbance.

We quantified stability using biodiversity metrics (species richness, community composition) rather than functional traits for two key reasons. First, biodiversity metrics capture aggregated biological activities that drive energy and material fluxes across spatiotemporal scales, serving as proxies to infer changes in ecosystem functions and services following disturbance (Snelgrove et al., 2014). Second, direct assessment of functional stability is hampered by fundamental ambiguities in defining ecosystem functions and the lack of standardized criteria for quantifying them (Garland et al., 2021). However, measuring changes in biological communities, especially in the ocean, presents challenges that prevent the use of traditional monitoring techniques: hydrodynamic forces (e.g., currents, waves) impede consistent sampling, while turbidity limits the reliability of visual surveys. Autonomous Reef Monitoring Structures (ARMS) offer standardized, high-resolution biodiversity assessments critical for reconstructing marine benthic communities in rapidly changing ecosystems. Studies using ARMS can reveal up to 1,200 operational taxonomic units (OTUs) per ARMS unit, spanning more than 30 phyla—a testament to their capacity to untangle intricate community structure under various anthropogenic impacts (McIlroy et al., 2024). ARMS were conventionally used as passive biodiversity samplers – deployed on the seabed for a certain period and then retrieved and processed to study the diversity that accumulated overtime (Leray & Knowlton, 2015; Obst et al., 2020). Here, we employed a novel, three phases experimental application of ARMS to assess the resistance and resilience of representative marine communities to anthropogenic stressors. In the first phase, ARMS were seeded with a baseline community in one of two marine protected areas (MPA) for twelve months (Figure 1a). In the second phase, ARMS with established baseline communities were translocated to areas of stress where they remained for twelve months to assess the response and resistance of that community to each human impact (Figure 1b). In the third phase, transplanted ARMS were returned to their seeding MPA site to evaluate community recovery and resilience (Figure 1c). We hypothesized that 1) human impacts would reduce species richness and restructure established communities; and 2) greater intensity impacts will result in stronger community responses eroding community resistance and resilience to stressors.

**Figure 1.**
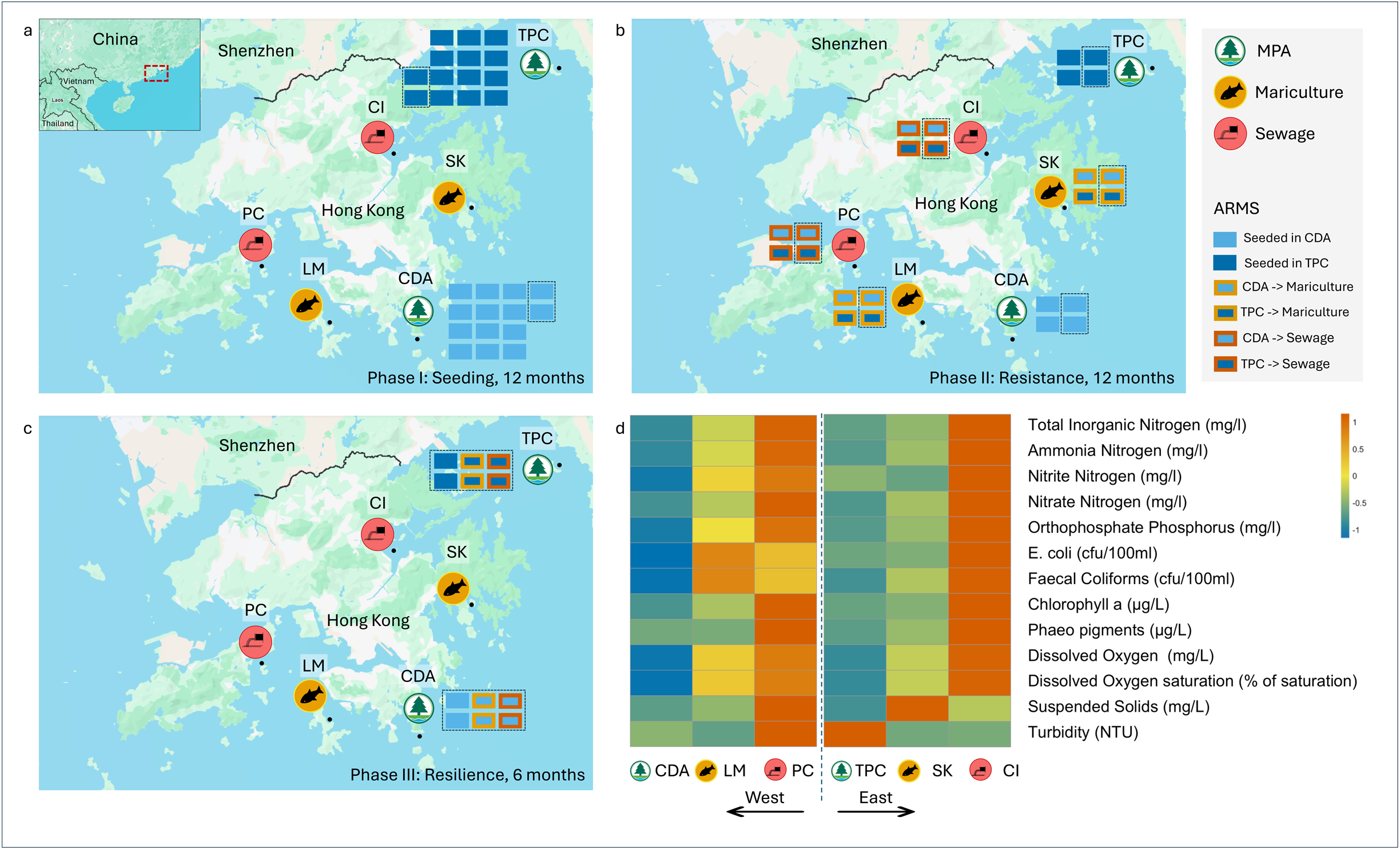
Six study sites in Hong Kong with three types (MPA : Cape D’Aguilar [CDA, west]/Tung Ping Chau [TPC, east]; mariculture : Lamma [LM, west]/Sai Kung [SK, east]; sewage : Peng Chau [PC, west]/Center Island [CI, east]) and their environmental impacts. (a) Phase I (Seeding): 28 ARMS deployed (14/site) in MPAs for 12 months; 2 ARMS/site (dashed outlines) sampled. (b) Phase II (resistance): 4 ARMS translocated from MPAs to impacted sites (mariculture: LM/SK, sewage: PC/CI). After 12 months, half of the ARMS (dashed outlines) were sampled. (c) Phase III (resilience): remaining ARMS returned to original MPAs; all ARMS sampled after 6 months. (d) Heatmap of water quality during Phase II, normalized by regional means (east/west). Red/blue indicate values above/below the mean (dissolved oxygen inverted). Map retrieved from Google Earth (Google, 2025). Map line delineates study areas and do not necessarily depict accepted country boundary.

## 2. Methodology

### 2.1 Study sites

The experiment was conducted in Hong Kong (22°18’00.0"N, 114°12’00.0"E) where relatively high biodiversity spans a complex environmental matrix. The city hosts over 6,000 described marine species representing 25% of China’s recorded marine fauna (Ng et al., 2017), with a recent ARMS-based study showing higher diversity in Hong Kong than Singapore, West Atlantic, and the Red Sea (McIlroy et al., 2024). In addition, there is a general water gradient where the west is heavily impacted by the Pearl River which discharges over 250 billion m^3^/yr of fresh water and about 22 million MT/yr of sediment from the mainland (Wei et al., 2020), and the east, exposed to the South China Sea, is more oceanic (Lai et al., 2016). Superimposed on this gradient are diverse anthropogenic stressors: The city is home to a population of over seven million with thriving residential, commercial, and industrial activities which generate over 2.8 million m^3^ wastewater daily. There are currently 65 wastewater treatment facilities across the city to process and disperse the effluent through submarine outfalls (HKDSD, 2019). Historically a fishing village, the city hosts 28 mariculture zones with about 910 licensed fish farms (HKAFCD, 2024a). Although operating on a small scale, these fish rafts impact water quality locally (Leung et al., 2015); Nonetheless, amidst these seemingly negative impacts, the government established nine marine protected areas (MPA) aiming to conserve and protect marine resources and conserve biodiversity (HKAFCD, 2024b). These mixed effects across the east-west gradient create diverse ecological niches leading to high beta-diversity (Stein et al., 2014). This regional diversity, coupled with a complex environmental gradient, makes Hong Kong an ideal place to study how coastal marine biodiversity changes under different levels of human impact. Six sites, representing 3 treatment types (marine park, mariculture, sewage) at either end of the east-west gradient, were chosen as ARMS deployment sites (Figure 1). They encompass two marine protected areas: Tung Ping Chau Marine Park (east) and Cape D’Aguilar Marine Reserve (west), two mariculture sites: Sai Kung Tai Tau Chau (east) and Lamma Lo Tik Wan (west), and two sewage sites: Center Island (east) and Peng Chau (west). Detailed site descriptions can be found in the supplementary materials (Section 1.1).

### 2.2 ARMS translocation

Autonomous Reef Monitoring Structures (ARMS) are standardized biodiversity samplers developed by NOAA and adopted by the Smithsonian Institution to investigate global marine biodiversity (NMNH, n.d.-a). ARMS consist of 9 stacked PVC plates (22.5 cm x 22.5 cm x 0.7 cm each) assembled with spacers between each plate that create small crevices for marine organisms to settle. Typically, they are deployed in the study area for a certain period then retrieved for processing. In this study, in order to test our hypotheses and understand how environmental factors influence marine communities, we applied a novel experimental approach consisting of three phases:

#### Phase I: Seeding Phase

In June 2020, we deployed 28 ARMS in the two MPAs: TPC and CDA (14 ARMS per site; Figure 1a). After a seeding period of 12 months, we processed 2 ARMS (termed baseline ARMS) from each site in June 2021 to characterize the initial community composition.

#### Phase II: Resistance Phase

At the end of phase I, we translocated 2 ARMS from TPC and CDA (4 total) to each of the four stressed sites (SK/LM, CI/PC). After the translocation, there were 4 ARMS at each of the 6 sites (24 ARMS in total; Figure 1b). These ARMS were exposed to the different disturbance treatments (fish farm, sewage outflow, negative control) for 12 months. In June 2022, we collected and processed 2 ARMS per site to assess community resistance (termed resistance ARMS). For the ARMS at the four stressed sites, we processed 1 ARMS seeded in TPC and 1 ARMS seeded in CDA (Figure 1b).

#### Phase III: Resilience Phase

After phase II, all ARMS at the stressed sites were translocated back to their original seeding site (e.g., ARMS moved from TPC to a stressed site were returned to TPC). By this point, 12 ARMS remained, all of which were deployed back to the MPAs (Figure 1c). In Dec 2022, we collected and processed all the remaining ARMS (resilience ARMS) to assess how the communities recovered from stresses. ARMS processing and molecular workflows followed established protocols (NMNH, n.d.-b; McIlroy et al., 2024). In brief, we first isolated three bulk fractions – two motile via sieveing (2 mm–500 µm, 500–106 µm) and one sessile by scrapping organic materials off the plates; then extracted DNA from these bulk fractions and amplified the mitochondrial COI region; lastly reconstructed community profile via metabarcoding for biodiversity analysis. Step-by-step protocols are detailed in supplementary materials (Section 1.2 ∼ 1.3).

### 2.3 Water quality

Water quality monitoring was conducted by the Hong Kong Environmental Protection Department (HKEPD). Over 20 parameters at three depths (surface: 1m below surface, middle: half-depth, and bottom: 1m above seabed) were measured monthly over 76 open water monitoring stations (HKEPD, 2024). We focused on 13 parameters (Table 1) that collectively represent major anthropogenic stressors and water quality gradients pertinent to our hypotheses. When available, we used data retrieved from the sampling station within our treatment sites; otherwise we calculated the mean value from several nearby sampling stations to reflect site water quality (Table S1). To visualize the data with a heatmap (Figure 1d), we calculated the annual mean of each of the 13 parameters during the period of deployment at treatment sites (resistance phase), normalized by standard deviation from the mean. Dissolved oxygen concentration values were inverted as higher values are considered a positive indicator of marine quality.

**Table 1.**
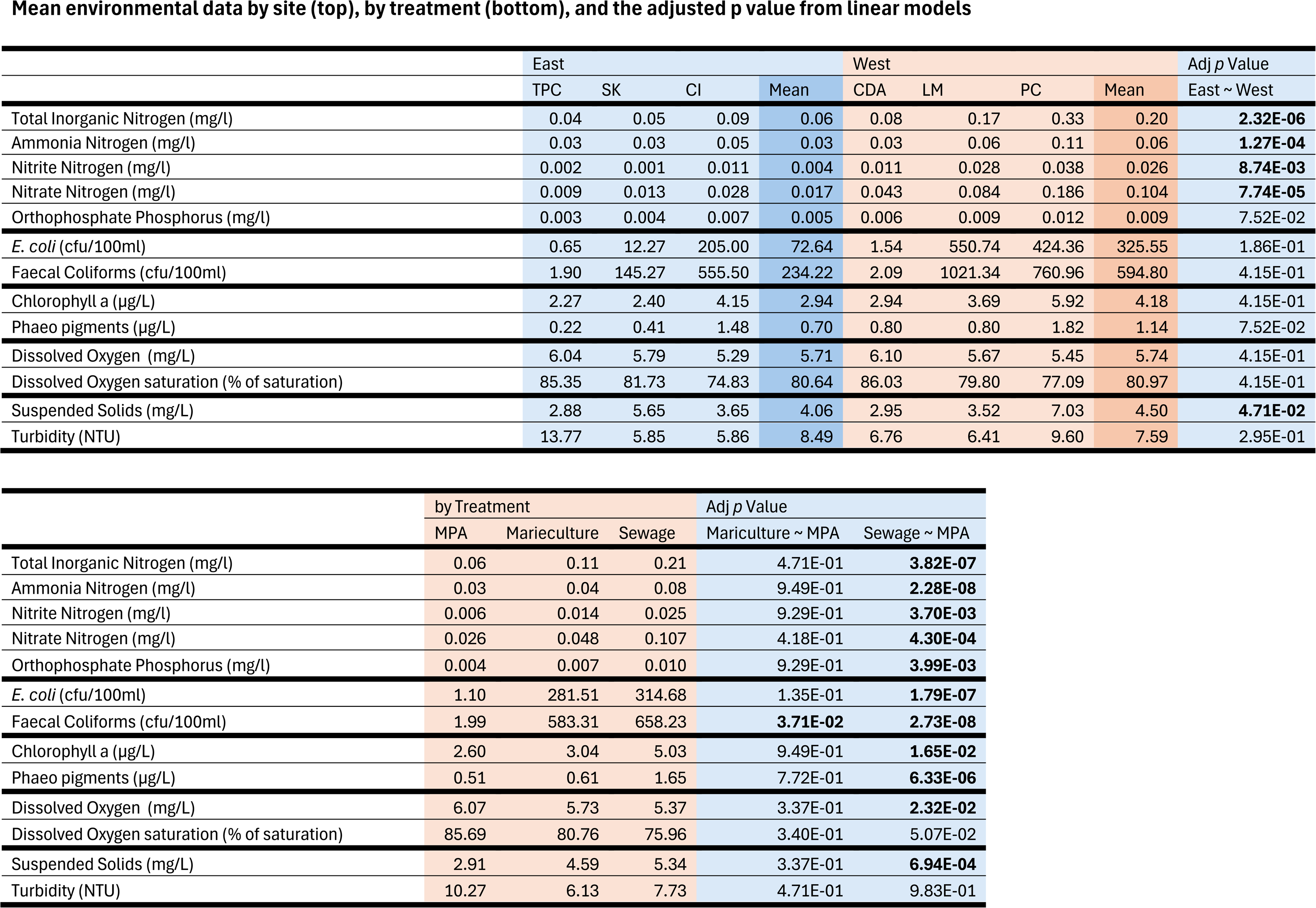
13 water quality parameters. Values within the East/West columns are annual mean value (June 2021-June 2022) of different sites, and values within the treatment columns are the annual mean value (June 2021-June 2022) of the three treatment types across east and west. *p*-values adjusted using the Benjamini-Hochberg procedure, with **bold numbers** indicating statistical significance (*p* < 0.05, all model df = 3,63).

We conducted a permutational multivariate analysis of variance (PERMANOVA, set.seed(100)) to test the effect of treatments and region (east vs. west) on water quality using monthly data of the 13 parameters from each site (adonis2 in the vegan package; Oksanen et al., 2007). To assess differences in water quality between the east and west regions and among treatments, we constructed a separate linear model for each environmental parameter using region and treatment as independent variables. Transformations were applied to skewed data of environmental metrics based on the distribution of the variables. To address left-skewed data, values were reflected by subtracting each observation from the dataset maximum, adding 1 to avoid zeros, and taking the natural log. For right-skewed data, a log transformation was applied directly. *p*-values were adjusted using the Benjamini-Hochberg procedure to account for the increased likelihood of false positives arising from multiple statistical tests (Benjamini & Hochberg, 1995).

### 2.4 Biodiversity analysis

Biodiversity analyses were conducted at the ARMS level, where data for the three fractions (motile 500µm, motile 106µm, and sessile) retrieved from the same ARMS were merged for analysis in Rstudio (2024.04.2 Build 764). Alpha diversity was measured by the number of OTUs (OTU richness) to facilitate comparison with other ARMS metabarcoding studies (Ip et al., 2023; Leray & Knowlton, 2015; McIlroy et al., 2024; Pearman et al., 2018). For the resistance ARMS, separate linear models were constructed to answer two key questions: 1) how do different treatments and water regions influence species richness? And 2) does species richness correlate with different environmental metrics? For non-normally distributed variables, transformations were applied following the same method outlined in section 2.3. A two-way ANOVA was conducted using treatment and water region as independent variables to examine whether they had an effect on 1) total OTU richness, and 2) richness within the six most abundant phyla. To assess successional changes in species richness, we modeled the total OTU richness and the richness of the six most abundant phyla within communities in the MPA over the study period. These were modeled as a function of deployment time (soak time) using Negative Binomial generalized linear models (GLMs) with a log link function. All *p*-values from multiple comparisons were adjusted using the Benjamini-Hochberg procedure

Beta diversity was analyzed with both Jaccard (qualitative) & Bray Curtis (quantitative) metrics after read counts were square root transformed, which mitigates extreme numbers contributed by PCR/sequencing bias (Van den Bulcke *et al*., 2023). To reveal how communities shift over treatments, we created ternary plots based on Jaccard metrics following a mathematical framework developed by Carvalho et al. (2012). Specifically, Jaccard similarity (β_sim_) is calculated with function (1)

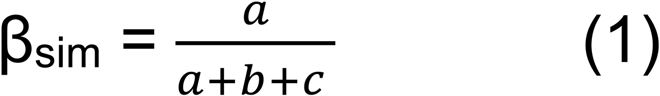

Where *a* is the number of shared species in both communities, *b* is the number of unique species of the first community, and c is the number of unique species of the second community. Then we defined two indices: community richness difference (β_rich_), measured by the different numbers of unique species between the two communities with function (2), and community replacement (β_rp_), calculated by the number of substituted species with function (3).

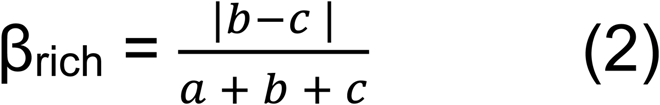

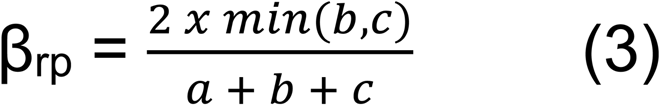

Ecologically, the differences between two communities can be explained by a combined effect of species richness difference and species replacement. Mathematically, the sum of the three indices is 100% (function 4), which allows us to visualize pairwise diversity in a ternary plot.

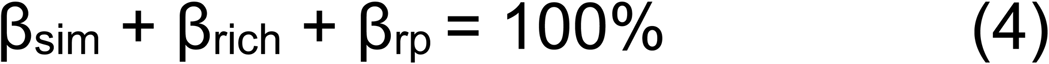

Two ternary plots were created, one for the resistance phase and the other for the resilience phase, to compare communities in sewage/mariculture treatment with those in the marine park of the same phase (Figure S2). Student’s t-test was used to identify whether the compositional change patterns were different between the two phases.

To understand if and how different communities clustered among treatments, a Bray Curtis distance matrix was first calculated (vegdist function in the vegan package). Principal coordinate analyses were conducted using the matrix over different treatments and visualized with ggplot2 (Wickham, 2016). Visualization was followed up by PERMANOVA (adonis2 in the vegan package, set.seed(123)) to reveal statistical differences among treatments across the different phases.

## 3. Results

### 3.1 Site characterization

The heatmap highlights the east-west water quality gradient and the severe impact in the sewage sites (Figure 1d). The west sites had significantly higher nutrient inputs (total inorganic nitrogen, ammonia, nitrite, and nitrate), and suspended solids than the east (Table 1); sites in the sewage treatment were heavily impacted with 10 water parameters significantly elevated and dissolved oxygen (DO) depleted relative to the MPA; mariculture sites showed significantly higher fecal coliforms than the MPA (Table 1).

Environmental conditions were significantly different among treatments (PERMANOVA, F_2,63_ = 3.51, *p* = 0.048, R² = 9.97%), but not between east and the west (F_1,63_ = 0.34, *p* = 0.69, R² = 0.48%). Pairwise comparisons revealed sewage sites differed substantially from MPAs (pairwise.adonis, *p* = 0.003, R² = 14.56%) while mariculture sites showed more moderate divergence (*p* = 0.027, R² = 11.20%).

### 3.2 Alpha diversity

Across all three phases, the 28 ARMS yielded a total of 16 million paired-end reads from three fractions (motile 500 µm, motile 106 µm, sessile), represented by 7,696 OTUs. Individually, the number of OTUs by ARMS varied from 704∼1,432 with a mean of 1,010. All 28 units harbored unique OTUs (50–217 per ARMS), with 3,463 OTUs (45.0% of total) occurring exclusively in single ARMS (Table S2).

For the resistance phase: 1) ARMS in the west hosted significantly lower OTU richness than the east (linear regression, t_3,8_ = 2.40, *p* = 0.043), and 2) ARMS from sewage sites hosted significantly lower richness compared to the baseline ones in the marine park (linear regression, t_3,8_ = 3.74, *p* = 0.0057). Analysis of environmental data (Table 2) revealed distinct categories of predictors for OTU richness based on their statistical significance and explanatory power:

1. **Strong predictors** (*p* < 0.05, R^2^ > 40%): ammonia nitrogen and total inorganic nitrogen (TIN) were identified as highly significant predictors of OTU richness, explaining over 40% of the variance.
2. **Moderate predictors** (*p* < 0.05, 40% > R^2^ > 20%): bacterial concentration, nitrate/nitrite nitrogen, and phaeopigment concentration showed moderate explanatory power.
3. **Weak predictors** (*p* < 0.05, R^2^ < 20%): orthophosphate phosphorus, suspended solids, dissolved oxygen (% saturation) had weak but statistically significant effects.
4. **Non-significant predictors** (*p* > 0.05): Dissolved oxygen concentration and turbidity were not significantly correlated with OTU richness.

**Table 2.**
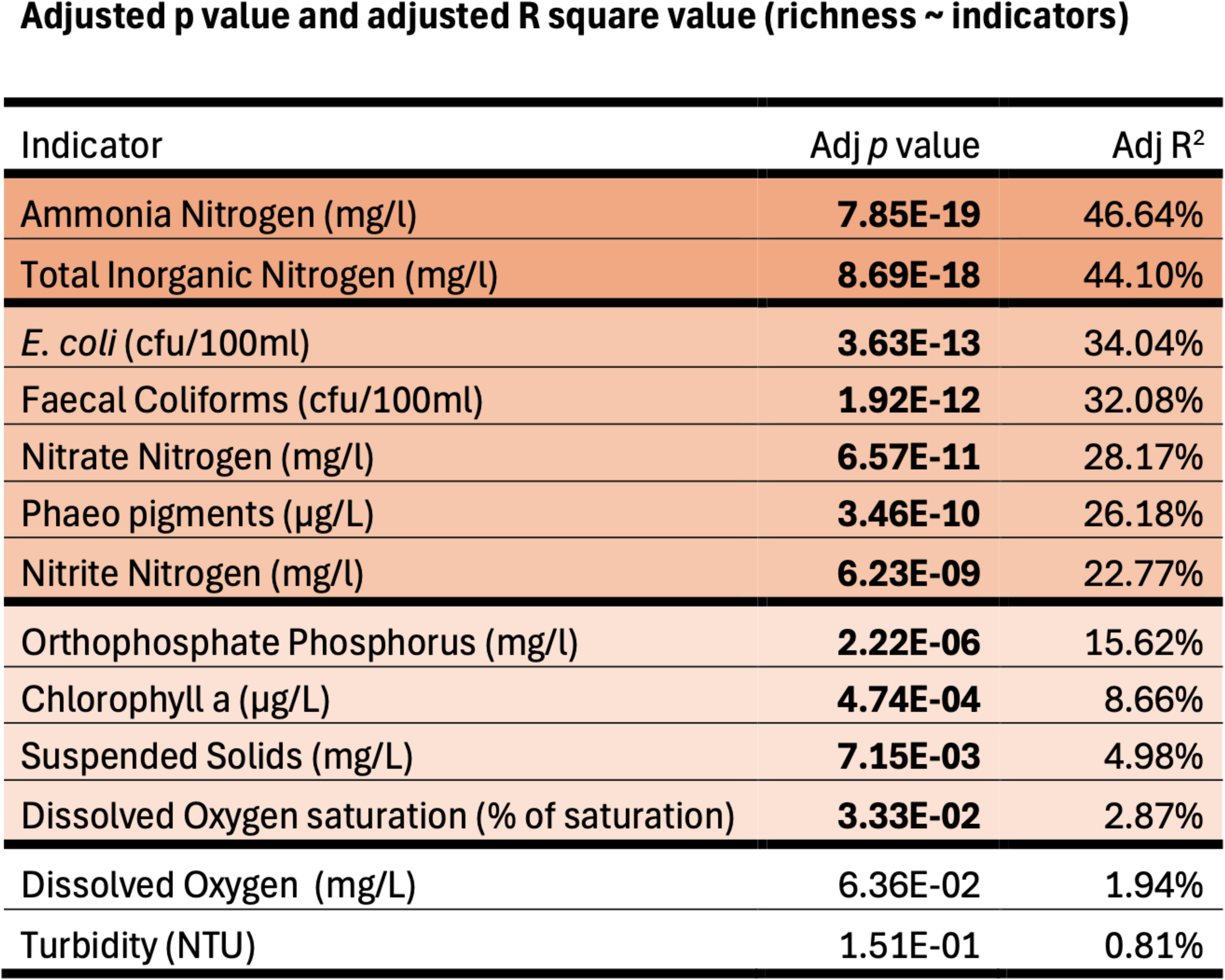
Benjamini-Hochberg adjusted *p*-values and adjusted R^2^ values from linear models (all models: df = 1, 132) testing associations between species richness and environmental metrics. All chemical indicators (ammonia nitrogen, TIN etc.), and bioindicators (faecal coliforms, phaeo Pigments) showed significant correlations with species richness.

These results highlighted the strong influence of nitrogen-related parameters on OTU richness, while dissolved oxygen, suspended solids, turbidity, and other non-significant predictors had little power to predict OTU richness. For the resilience ARMS, the one-way ANOVA revealed no significant effects of treatments on OTU richness (f_2,9_ = 2.83, *p* = 0.11), suggesting that OTU richness in stressed communities returned to similar levels following a 6-month recovery.

Two ARMS were sampled in each of the MPA by the end of every phase. We found OTU richness declined in these MPA communities over the study period (Figure 2a). More specifically, from the OTU counts retrieved from the MPA ARMS, the mean OTUs decreased from 1327 (seeding phase, 12 months) to 1205 (resistance phase, 24 months), and then 894 (resilience phase, 30 months). GLM models revealed a steady decrease in species richness over the succession process (glm.nb, west communities: z = −3.11, *p* = 0.002, east communities: z = −3.677, *p* = 0.0002. Table S3, Figure 2a).

**Figure 2.**
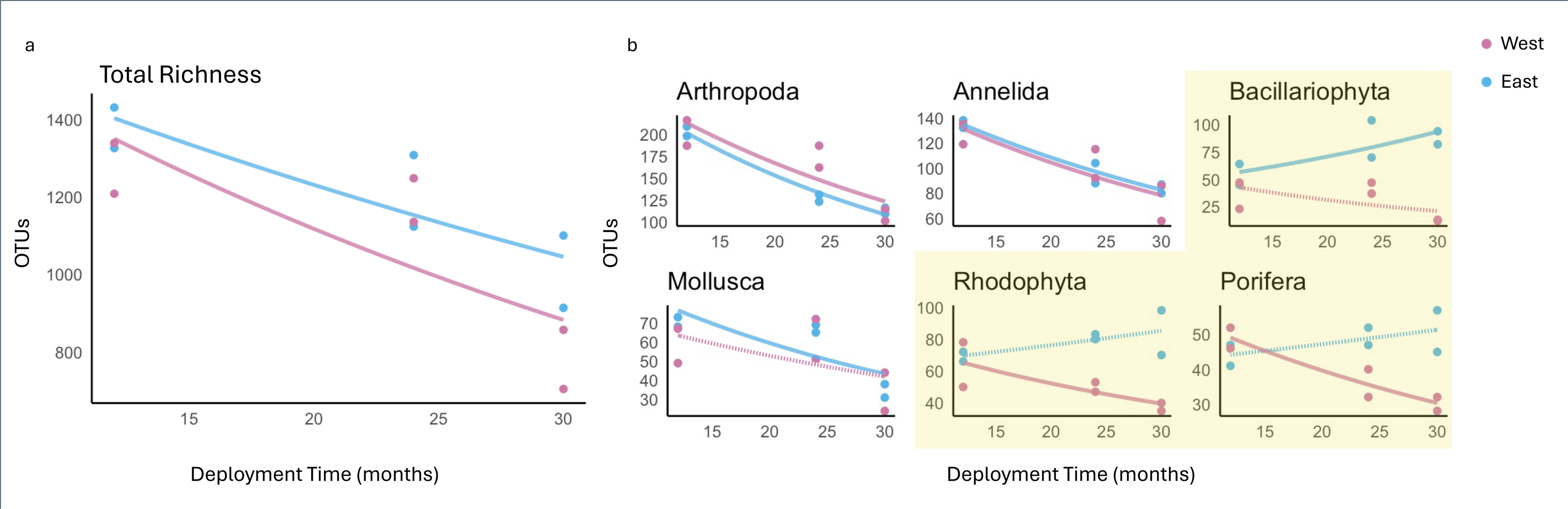
(a) OTUs in east (TPC, blue) and west (CDA, purple) MPAs over three phases: seeding (12 months), resistance (24 months), and resilience (30 months). Richness declines progressively in both MPAs. (b) Phylum-level richness trajectories for the six most abundant taxa. Sessile taxa (Bacillariophyta, Rhodophyta, Porifera; yellow boxes) exhibit different successional patterns between MPAs: East communities (TPC) sustain stable or increasing richness, while west (CDA) declines. All trends modeled with Negative Binomial regression. Dash lines indicated the log-rate slope was not statistically significant (p>0.05).

### 3.3 Beta diversity

From the resistance ARMS, we found high species turnover among different treatments (Figure S2). Using the Jaccard metric for comparisons, communities from the stressed sites and the marine parks differed by 84.2%, where 73.3% of the difference was explained by species turnover, and 10.9% was explained by richness difference. Resilience ARMS that were translocated back to the two MPAs exhibited a different pattern. The same comparison showed that communities were 66.2% different, where 52.6% of the difference was explained by species turnover, and 13.6% explained by richness difference. Student’s t-tests revealed significant differences between the changes in compositional difference (t_30_ = 7.16, *p* < 0.001) and turnover (t_30_ = 5.56, *p* < 0.001) but no significant changes in richness differences (t_30_ = 1.02, *p* = 0.318), which indicated significantly different compositional changes contributed by higher species turnover in the resistance phase.

PCoA plots based on Bray-Curtis metric showed clear clustering among different treatments. There were clear spatial and temporal clusters of ARMS recovered from the MPAs (blue, Figure 3) The PERMANOVA test confirmed significant effects of deployment time (F_2,6_ = 9.67, *p* = 0.002), water region (F_1,6_ = 44.67, *p* = 0.001), and their interaction (F_2,6_ = 9.16, *p* = 0.001) on community composition. All together, they explained 93.21% of the observed variation. The post hoc Pairwise PERMANOVA revealed significant differences between all phase pairs (seeding - resistance: F_1,4_ = 8.04, *p* = 0.017; seeding - resilience: F_1,4_ = 16.84, *p* = 0.007; resistance - resilience: F_1,4_ = 4.76, *p* = 0.035), with water region (east/west) consistently explaining the most variation (seeding - resistance: R^2^ = 47.36%, F_1,4_ = 20.74, *p* = 0.004; seeding - resilience: R^2^ = 41.50%, F_1,4_ = 24.56, *p* = 0.004; resistance - resilience: R^2^ = 81.17%, F_1,4_ = 51.72, *p* = 0.006).

**Figure 3.**
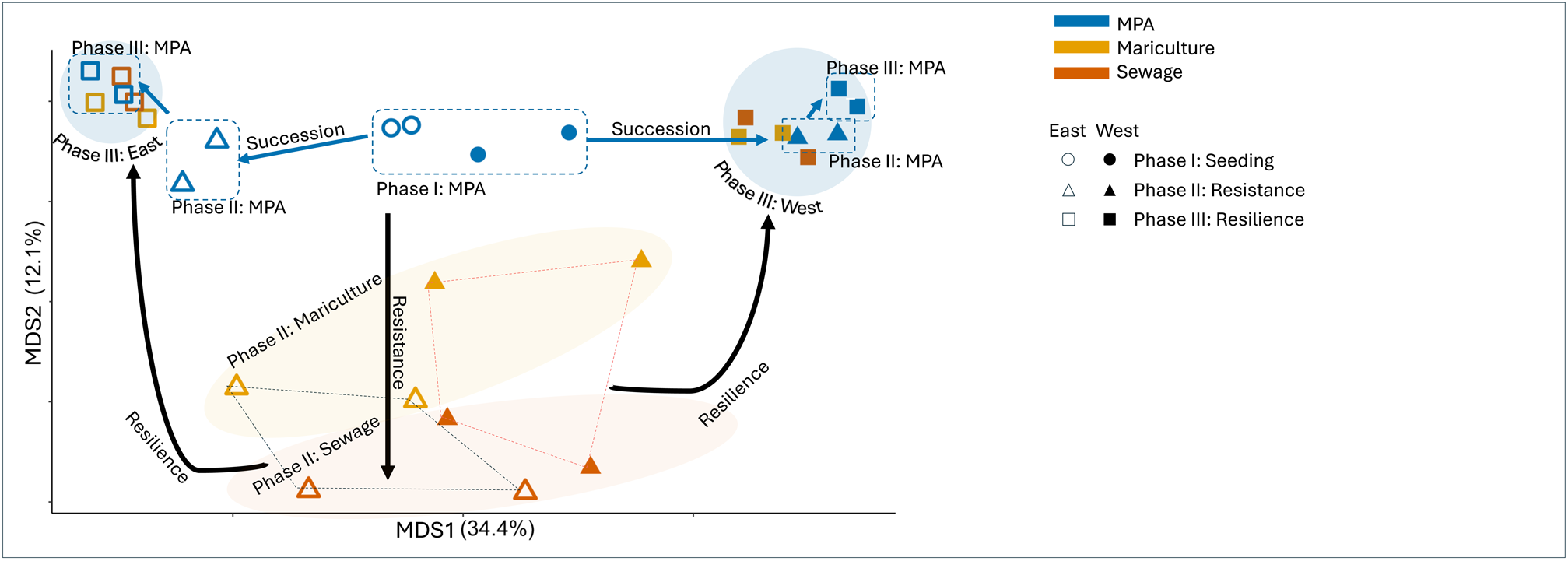
PCoA of all ARMS (Bray-Curits metric). Each point represents one community in different sites (MPA: blue, mariculture: yellow, sewage: red) from different phase (phase I: circle, phase II: triangle, phase III: square) across the east (hollow shape) and the west (solid shape). **Succession** was indicated by blue shapes in the top part of the plot where it started in the middle (circles) in the seeding phase, developing sideward through the resistance phase (triangles), and reached to both top corners (squares); communities translocated to the stressed sites during **resistance** phase moved downward with sewage communities on the bottom; after all communities were translocated back to the two MPAs in the **resilience** phase, all communities from the same seeding site clustered together on the two top corners. Note that in the west, phase II MPA communities (green triangles) were clustered together with all the resilience communities (top right) but the pattern is different in the east (top left).

The resistance ARMS (triangles, Figure 3) demonstrated strong compositional control by treatment, with baseline communities clustering in the upper quadrant, mariculture communities intermediate, and sewage communities in the lower part (PERMANOVA: F_2,6_ = 4.15, *p* = 0.002, R² = 24.1%). This stratification reflected a gradient of human impacts from baseline (least impacted) to sewage (most impacted). The regional water quality gradient further structured communities, with eastern sites clustering in the left and western sites in the right (PERMANOVA : F_1,6_ = 10.12, *p* = 0.001, R² = 29.4%). The significant treatment – regional interaction (PERMANOVA: F_2,6_ = 5.01, *p* = 0.001, R² = 29.1%) indicated that treatment responses varied regionally, suggesting the strength of local disturbance was impacted by broader environmental gradients.

For resilience communities (squares, Figure 3), the TPC/CDA communities were closely clustered on the top left/right side of the quadrant respectively regardless of pre-exposed disturbance, indicating the significant compositional changes in the resistance phase were no longer evident after the resilience phase (PERMANOVA: site: F_1,11_ = 54.32, *p* = 0.001, R² = 84.2%; treatment: F_2,11_ = 1.09, *p* = 0.25, R² = 0.03%). Additionally, the TPC (east) communities (hollow squares) were tightly clustered without overlapping the MPA community in the resistance phase (hollow triangles), while the CDA (west) communities (solid squares) were loosely clustered overlapping the MPA community in the resistance phase (solid triangles), indicating different recovery patterns between the two sites (PERMANOVA: phase x sites: F_1,12_ = 3.41, p = 0.05).

### 3.4 Taxonomic assignment

Of the 7696 clustered OTUs, 4009 (52.1%) were assigned to the phylum level with a minimum of 80% similarity on at least 300 bp, and 856 (11.1%) were assigned to the species level with a minimum 95% similarity on at least 300 bp. The six phyla with the greatest detected species richness were Arthropoda, Annelida, Bacillariophyta, Rhodophyta, Mollusca, and Porifera. During the resistance phase, each of these phyla exhibited different responses across eastern and western sites (Figure 4b). We found higher richness of Arthropoda and Annelida in the heavier impacted west, while richness of Bacillariophyta, Rhodophyta, Mollusca, and Porifera were higher in the east.

**Figure 4.**
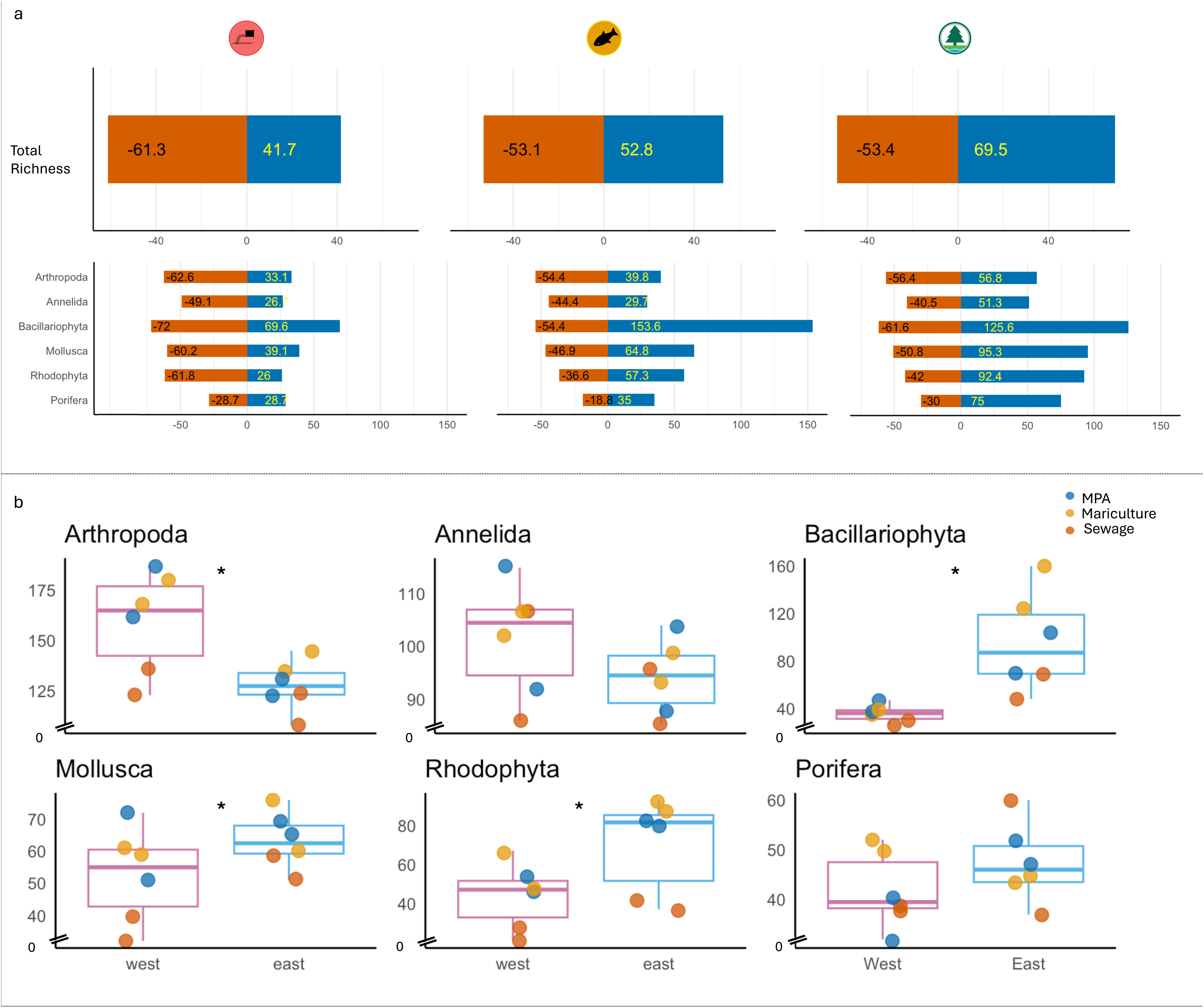
Changes in alpha diversity after resistance phase. (a) percentage total species richness loss (red) and gain (blue) of the resistance communities over the seeding communities (top); percentage species richness of the top six abundant phyla loss (red) and gain (blue) of the resistance communities over the baseline communities (bottom). (b) species richness difference over east and west by phyla. Asterisk indicate significant differences between east and west.

To better understand these compositional changes, total species richness and species richness of the major phyla from the resistance ARMS (2 treatment sites [mariculture, sewage] and the marine park) were compared with their respective baseline communities in the seeding phase. Net losses on OTU richness were found in both stressed treatments (19.6% in sewage and 0.3% in mariculture), while a 16.0% OTU richness increase was identified in the resistance phase MPA over the seeding phase baseline. By the end of the resistance phase, species richness of the top six phyla in the sewage treatment suffered the highest percentage loss, followed by mariculture where both net gains/losses were observed. In contrast, species richness increased in all top six phyla of the marine park community (Figure 4a).

Communities in the MPA ARMS over the three phases revealed different temporal richness changes in different taxonomic groups. The total OTU counts for the top six taxa that were retrieved from the west marine park (CDA) showed a gradual decrease over the three phases, mirroring the temporal changes in total richness. A similar pattern was found in the east MPA (TPC) with Arthropoda, Annelida, and Mollusca, but not with the sessile Bacillariophyta, Rhodophyta, and Porifera, suggesting different succession patterns (Figure 2b).

## 4. Discussion

Global biodiversity loss and shifts in community composition caused by anthropogenic stressors have been well documented since the late 1980s (Wilson, 1989). With expanding coastal populations, urbanization is escalating these impacts, particularly in coastal marine ecosystems (Sahavacharin et al., 2022). Using ARMS coupled with metabarcoding techniques, we investigated highly reproductive, yet short-lived, cryptic organisms – sensitive to environmental changes but are often overlooked by conventional macrofauna surveys that rely on visual observation. This approach also enabled community-level reconstruction of benthic diversity, revealing patterns that would have otherwise been undetectable through traditional methods – which are restricted to much smaller numbers of species. Our work found that sewage and mariculture, can drive major changes to community structure. However, significant recovery was evident following stress removal. Further investigation into the specific environmental traits that drove ecosystem instability identified nitrogen, particularly ammonia nitrogen, as the leading predictor of reductions in species richness and changes in community composition. Interestingly, communities showed signs of rapid recovery within six months, becoming more similar to the ARMS that remained in MPAs and were not exposed to stress. Low resistance, yet high resilience, were consistent features of these communities, likely underpinned by the influence of nearby oceanic waters which dilute impacts and facilitate high connectivity within marine environments. These experimental findings demonstrated how urban development impacts coastal marine biodiversity and provide critical insights for future conservation efforts to mitigate biodiversity loss.

### 4.1 Community resistance

Mariculture and domestic sewage are two umbrella terms that encompass a broad range of stressors including changes in habitats, excessive nutrient inputs, pathogen and heavy metal pollution, increased suspended solids, among others. Additionally, these stressors can interact synergistically and exacerbate negative outcomes (Drechsel et al., 2015; Martinez-Porchas & Martinez-Cordova, 2012; Wear & Thurber, 2015). While many of the environmental stressors overlapped among treatments, their intensity varied widely. Monthly environmental data indicated that sewage sites showed a greater degree of impact compared to the mariculture sites (Table 1, Figure 1d). Together with the diversity data, our findings revealed that both alpha (OTU richness) and beta (compositional shifts) diversity were shaped by regional water quality gradients and local anthropogenic stressors. Particularly, TIN acted as a primary predictor of biodiversity loss and it mediated treatment-driven restructuring of communities. These results align with global evidence linking eutrophication and its cascading impacts to the ecosystem. Many marine primary producers are nitrogen limited, including various species of phytoplankton, microalgae, and macroalgae, and elevated levels of TIN promote population growth in these species. Under moderate TIN enrichment, these favored species escalate space and light competition which negatively impacts a wide range of sessile taxa, decreasing species richness and changing the community composition. At severe TIN concentrations, the excessive growth of phytoplankton and algae can cause harmful algal blooms (HAB), which lead to hypoxia and, in extreme cases, ecosystem collapse (de Vries, 2021; Galloway et al., 2003; Wurtsbaugh et al., 2019). Additionally, nitrate and phosphate in high concentrations augment the photosynthesis-respiration cycle which increases pH variability, hindering calcification of reef building corals (Silbiger et al., 2018). Elevated reactive nitrogen such as ammonia, nitrite, and nitrate all show different levels of toxicity to marine life, impacting their survival, growth, and reproduction (Camargo & Alonso, 2006).

Species richness responses to anthropogenic stress varied across phyla: while total OTU richness declined with elevated nitrogen and bacterial indicators (Table 2), Arthropoda and Annelida richness were higher in heavily impacted western communities compared to eastern sites (Fig. 4b), revealing taxon-specific heterogeneity. In addition, we found high species turnover in all communities under stress (Figure S2) coupled with high percentages of ARMS-specific species (Table S2). Together, these results indicated a nutrient-driven restructuring of communities rather than a uniform biodiversity loss. Communities in highly impacted sites were not merely a subset of those in the pristine sites. Instead, heavy nutrient pollution profoundly affected the species make up of communities, inducing phase shifts towards an alternate state (deYoung et al., 2008; Lesser, 2021).

The environmental impacts driven by sewage and mariculture were correlated with significant shifts in both species richness and community composition, which underscored the low community resistance of urban coastal marine ecosystems to human-induced stressors, particularly nutrient pollution.

### 4.2 Community resilience

In this study, benthic communities exhibited considerable resilience. Significant changes in species richness and the community structure, induced by different treatments over the resistance phase (12 months), were reversed in the resilience phase (6 months). However, differences in recovery patterns were identified between the east and west. Communities seeded in TPC (east) demonstrated a more complete recovery, as indicated by tighter and more distinct clustering, whereas communities in the CDA (west) showed less defined clustering (squares, Figure 3). This pattern revealed subtle, but significant, site specific differences in community resilience.

Community resilience is centered around changes in community composition that are dictated by the species pool (Cornell & Harrison, 2014) and influenced by a series of complex factors, such as priority effect (Weidlich et al., 2021), dispersal (Álvarez-Noriega et al., 2020), environmental filtering (Kraft et al., 2015), and disturbance regime (Pirotta et al., 2018). Above all, a rich species pool is critical for enhancing community resilience because a diverse regional community is more efficient in utilizing resources and thus more productive (Wagg et al., 2022). Greater species richness also strengthens functional redundancy, where the loss (or failure in recruitment) of some species can be compensated by others (Fetzer et al., 2015), and it diversifies trophic interactions so that the food web is more robust (Thébault & Loreau, 2005). Furthermore, it provides a more complex community structure to promote species recruitment (Hooper et al., 2005).

The high biodiversity of our study region (Ng et al., 2017) may thus have contributed to high community resilience, aligning with ecological theory linking regional species pools to recovery capacity. Communities in CDA (west) – exposed to higher chronic nutrient loading than TPC (east, Table S4) – exhibited slower recovery, suggesting that, although regional biodiversity provided a reservoir for recovery which buffered against disturbance, elevated local stressors (e.g., TIN, orthophosphate) can impede these stabilizing effects. This finding emphasized the need for conservation strategies that couple regional biodiversity protection with targeted mitigation of local stressors to sustain resilience in rapidly urbanizing coastal marine ecosystems.

### 4.3 Succession pattern

While translocating communities to sites with different environmental conditions revealed their resistance and resilience to stress, extended monitoring uncovered succession-driven compositional shifts over time. Communities from ARMS in the two MPAs, which remained at the same sites throughout the study, steadily decreased in species richness from the 12 months, to the following sampling points at 24 months and 30 months (Figure 2a).

Initial colonization of newly available space begins with macromolecules and bacteria, followed by microalgae and larvae, and then macro-organisms (Noël et al., 2009). The pioneer colonizers promote the species richness and compositional diversity of the community revealed by the high OTUs at the end of the seeding phase ARMS. Over time, succession further increases both abundance and biomass, either through the success of dominant species, which reduces species richness via competition, or the coexistence of species leading to stable/increased species richness (Vicente et al., 2021). Our results revealed both taxon- and site-specific succession trajectories. Richness of Arthropoda, Annelida and Mollusca decreased temporally in both MPAs. Site dependent responses were exhibited in three primarily sessile taxa: Bacillariophyta, Rhodophyta, and Porifera. In CDA (west), these taxa mirrored the overarching decline, but in TPC (east), their richness increased during the resistance phase and plateaued at high levels in resilience phase (Figure 2b).

Both Bacillariophyta (diatoms) and Rhodophyta (red algae) are primary producers that share a competitive niche with Chlorophyta (green algae), Ochrophyta (brown algae), and photosynthetic Pyrrophyta (dinoflagellates). Together, they compete for nutrients, light, and space. Although CDA (west) and TPC (east) are both MPAs, mean TIN in CDA was over 90% higher than TPC due to the east-west gradient in water quality (Table S4). This higher nutrient exposure, particularly from nitrogen, favors fast-growing, bloom-forming species – specifically Bacillariophyta, Ochrophyta, and Chlorophyta (HKAFCD, 2023; Porter et al., 2013). The dominance of these opportunistic species likely displaced slow-growth algae and smothered sessile invertebrates like Porifera (sponges) which rely on open benthic space for recruitment and growth, ultimately reducing benthic diversity (Easson et al., 2014; Teichberg et al., 2010). Consequently, total richness of Bacillariophyta, Rhodophyta, and Porifera in CDA declined progressively to a low-diversity equilibrium, reflecting accelerated competitive exclusion under stable, nutrient-rich conditions. In contrast, TPC’s low-nitrogen environment facilitated coexistence among the sessile community with different growth strategies, buffering competitive exclusion and extending the high-richness pre-equilibrium phase. This divergence suggested a fundamental trade-off: faster community stabilization comes at the cost of reduced biodiversity.

## 5. Conclusion

By leveraging metabarcoding techniques, we expanded our understanding of community diversity changes in an urban megacity. Elevated human impacts, indicated by high TIN and bacterial concentrations, reduced species richness and undermined community resistance and resilience. Marine coastal communities were not resistant to acute human impacts but exhibited high community resilience. In addition, temporal sampling from the same sites revealed that nutrient pollution exerted phylum-specific effects, altering successional trajectories. These findings provide important insights for directing future conservation efforts: 1) alleviate local impacts – with a particular focus on nitrogen sources – as they are disproportionately disruptive to benthic diversity and community composition; 2) monitor regional biodiversity as it is a backbone supporting high community resilience. When integrated with high-resolution environmental monitoring, ARMS provide a framework to track biodiversity responses across anthropogenic stress gradients, empowering evidence-based conservation in human-altered seascapes

## 6. Data availability

Raw sequence data are openly available on figshare at https://doi.org/10.6084/m9.figshare.29481053. The rest of the dataset for analysis are available via github at https://github.com/zhongyuewan/MGEXP1.

## 7. Code availability

Data analyses were performed on RStudio (2024.09.0 Build 375). Codes are available via github at https://github.com/zhongyuewan/MGEXP1.

## 8. Acknowledgment

This research was supported by funding from the Research Grants Council, the University Grants Committee (研究資助局) (CRFC7013--19G). Fieldwork and ARMS processing were supported by our technician So Wing Wa Oscar, staffs of Swire Institute of Marine Science, teams of secondary school and university student volunteers. DNA sequencing supported by the Genomics Core Centre for PanorOmics Science (CPOS) of the University of Hong Kong.

## 9. Author Contributions

D.M.B. and S.E.M. conceived and designed the research. All authors undertook fieldworks and AMRS processing. Z.W., W.Y.Z., and C.H carried out DNA extraction. Z.W. analysed the data and wrote the first draft. All authors were involved in reviewing the article.

---

**Figure S1.**
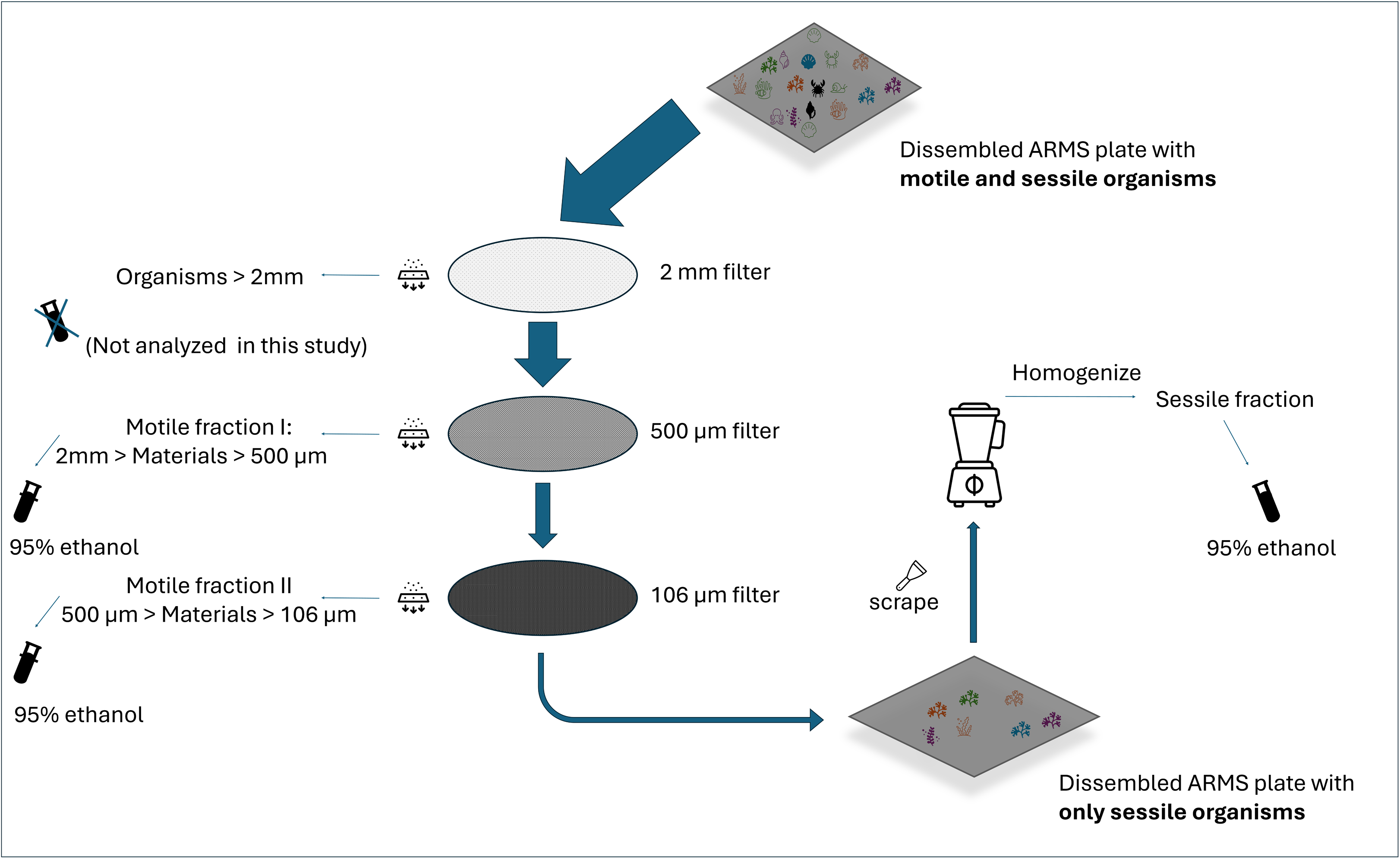
Post-dissembling, individual ARMS plates (top) were rinsed with filtered sea water to remove any motile organism and remaining debris. Dislodged materials were then sieved through a filter set of three (2 mm, 500 µm, 106 µm). Organisms over 2mm were not analyzed in this work. Motile fraction I and II were then preserved in 90% ethanol for later downstream processing. Sessile organisms retained on the plate were then scraped off and put though a blender to homogenize (sessile fraction), and then preserved with 95% ethanol.

**Figure S2.**
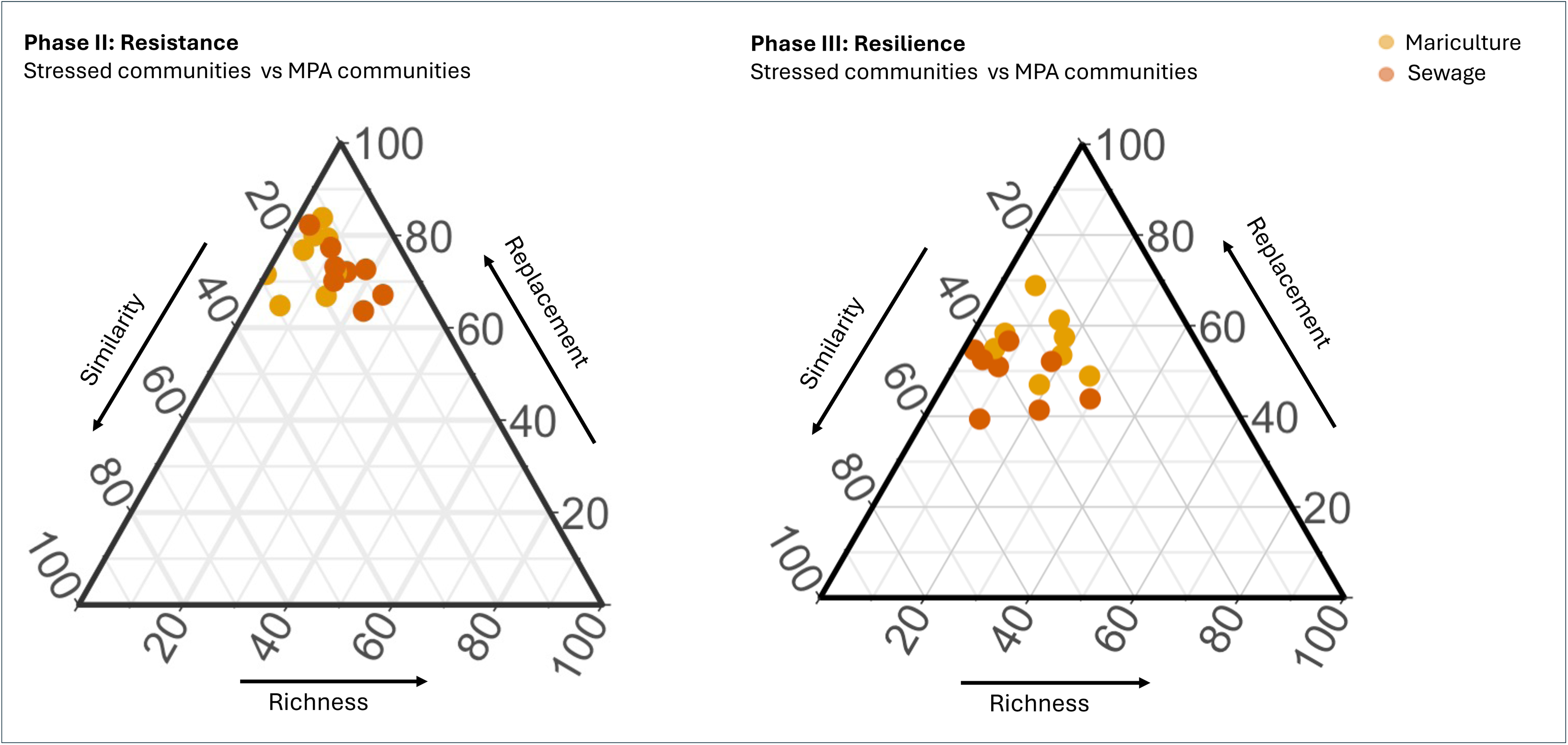
Ternary plots of beta diversity components (species replacement [βrp], richness difference [βrich]) during resistance (left) and resilience right) between stressed communities and MPA communities.

**Table S1.**
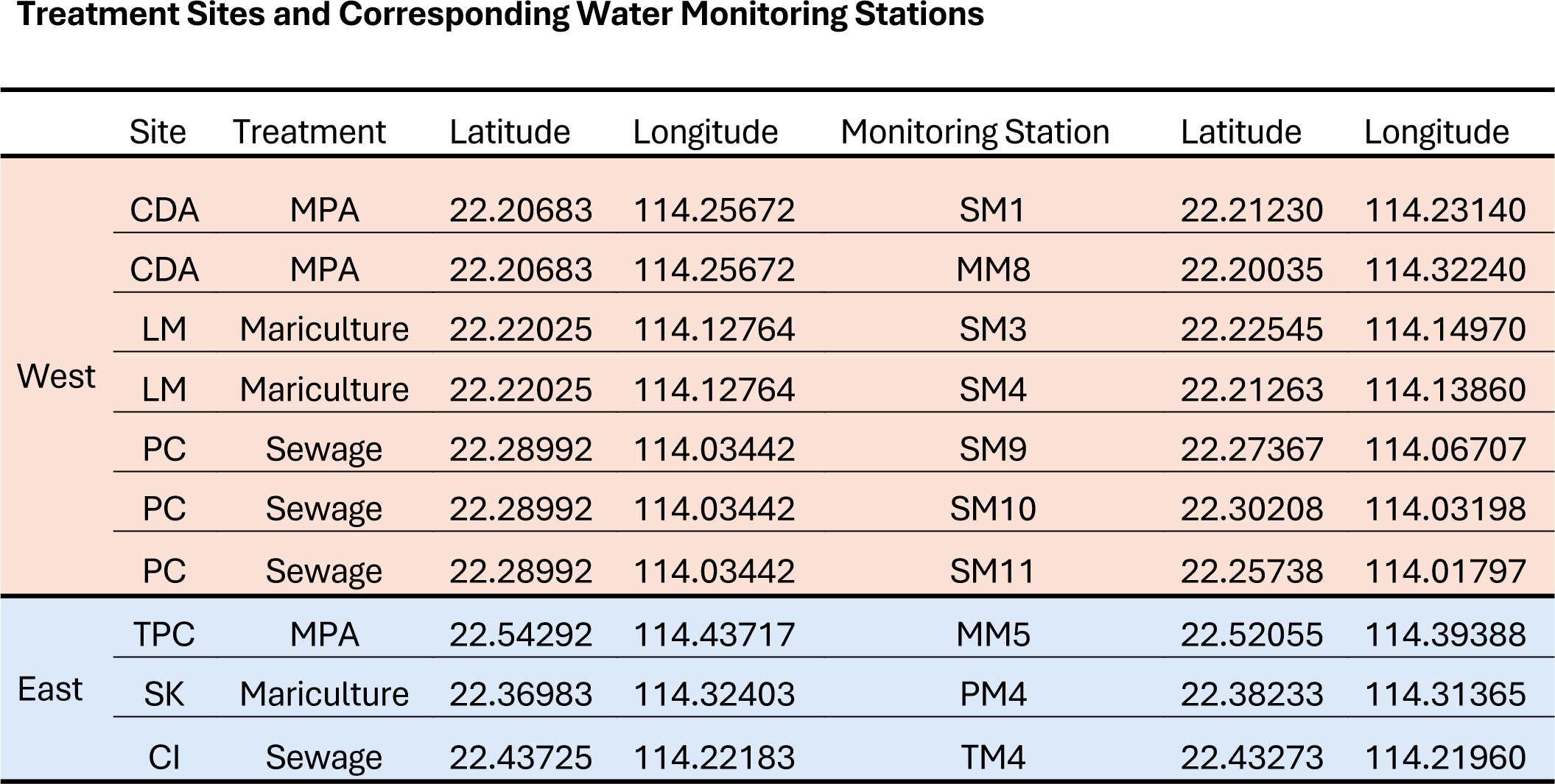
Treatment sites, their corresponding water monitoring stations and the GPS coordinates.

**Table S2.**
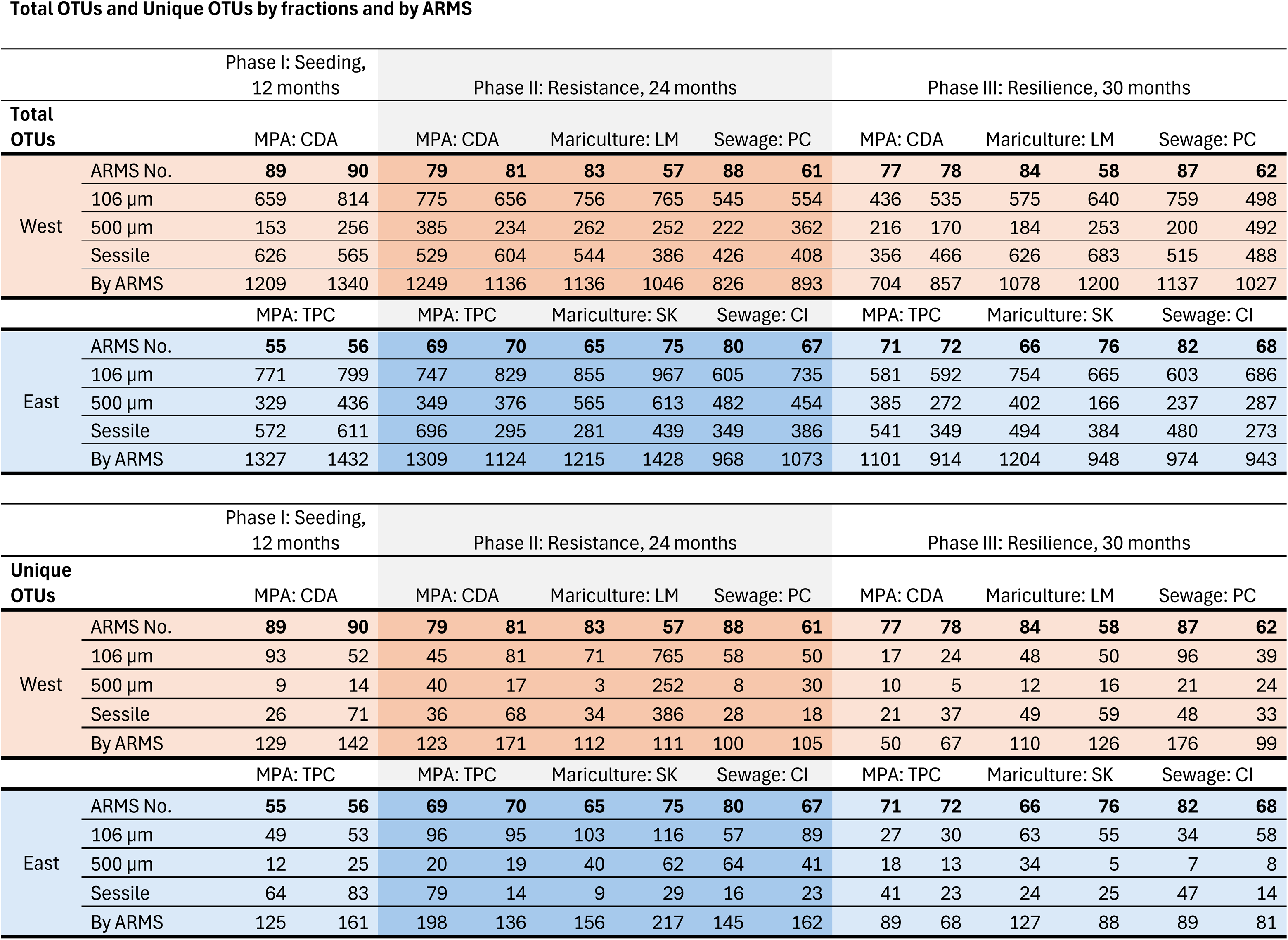
Total species richness (top) and unique species (bottom) by fractions and by ARMS.

**Table S3.**
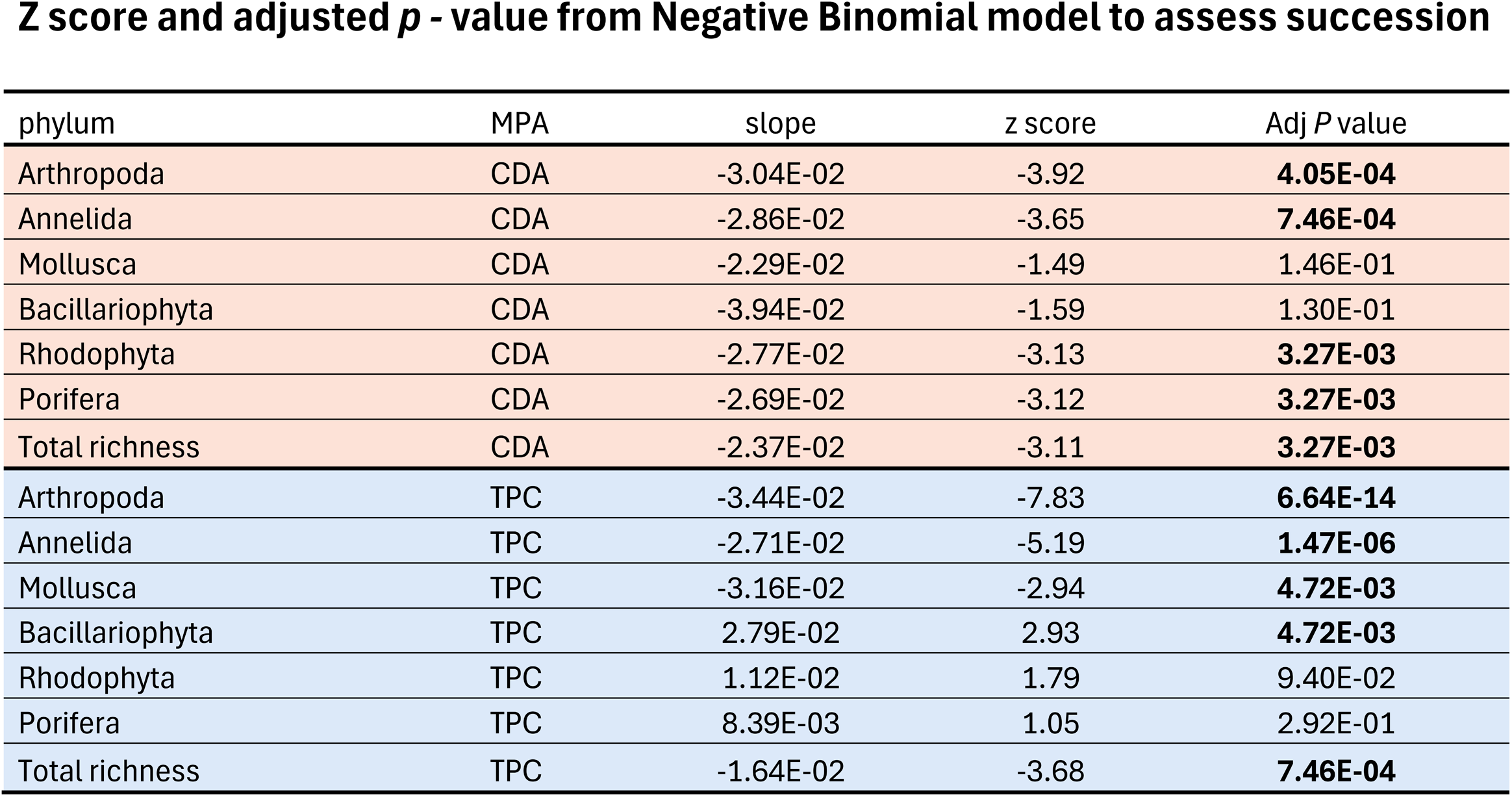
Z-scores and adjusted *p* values from Negative Binomial model to study community succession. Bold value highlighted in adjusted *p* indicated significant trends (*p* < 0.05 after Benjamini-Hochberg adjustment).

**Table S4.**
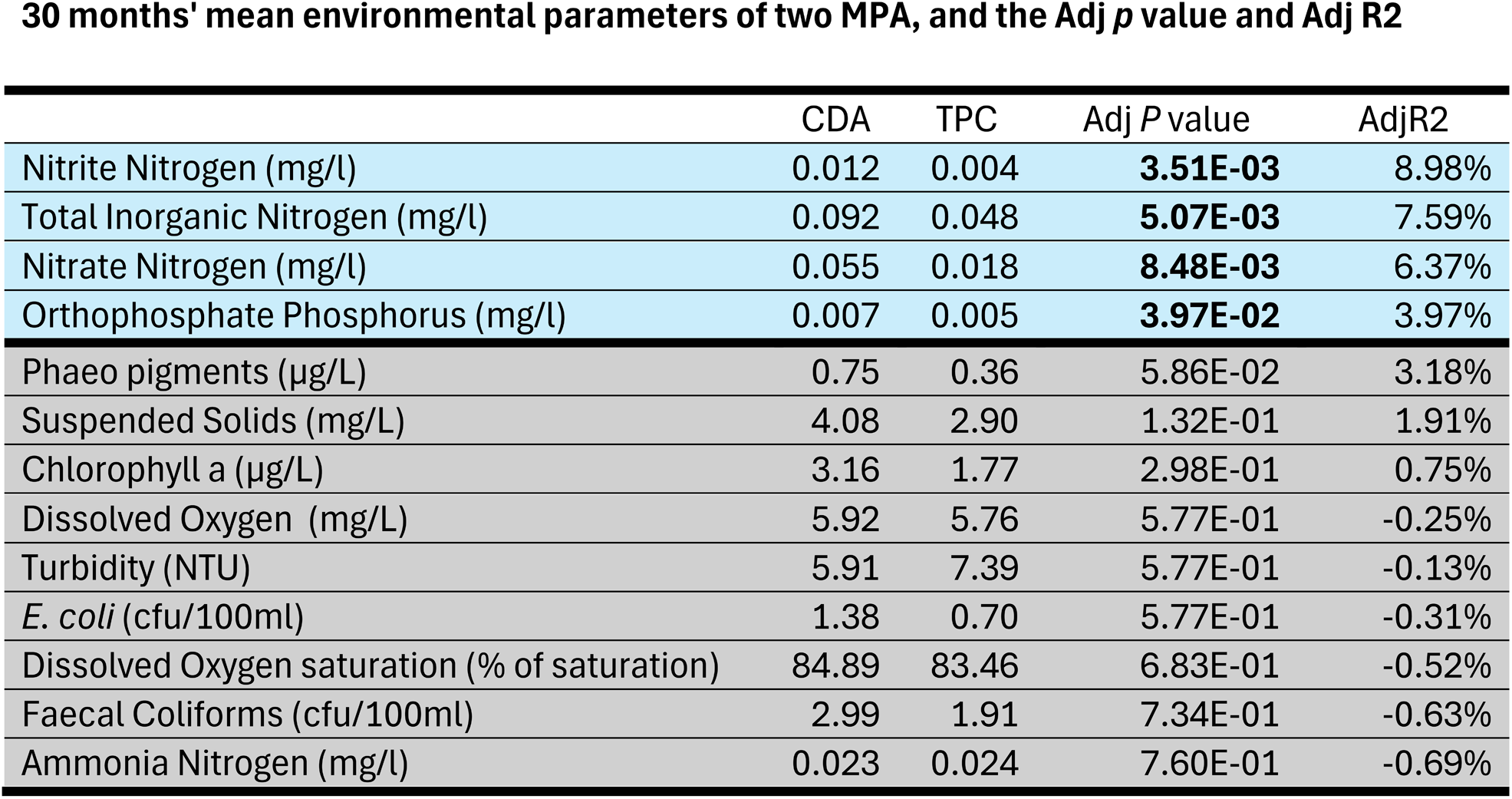
Mean values of all 13 environmental parameters over the study period (Jun 20 ∼ Dec 22) from two marine park (CDA, TPC), and adjusted *p*, adjusted R^2^ value from linear model (all df = 1,131). Parameters showed significant site differences were highlighted in blue (*p* < 0.05 after Benjamini-Hochberg adjustment).

