## Supplementary Materials for "Autonomous Reef Monitoring Structures (ARMS) Reveal Human-Induced Biodiversity Shifts in Urban Coastal Ecosystems"

Zhongyue Wan *et al.*

**The PDF file includes:**

- Materials and Methods
- Supplementary Figures
- Supplementary Tables
- References

### 1. Materials and Methods

#### 1.1 Site Description

Marine Park:

Tung Ping Chau Marine Park (TPC, 22°32'34.5"N 114°26'13.8"E) is a 270 ha MPA surrounding the northeasternmost island of Hong Kong (HKAFCD, 2024b). Sheltered by Mirs Bay and exposed to the oceanic South China Sea, it hosts a robust coral community of over 10 genera of scleractinian coral (Yeung et al., 2021). Cape D'Aguilar Marine Reserve (CDA, 22°12'24.6"N 114°15'24.2"E) is a much smaller MPA of only 20 ha located on the southeast tip of Hong Kong Island. Compared to TPC, it has a higher anthropogenic footprint due to the proximity to urban areas. Regardless, it is rich in biodiversity with different habitats including rocky and cobble beaches, rocky shore, intertidal pools, and coral communities (Xu et al., 2015).

Mariculture:

Sai Kung Tai Tau Chau (SK, 22°22'11.4"N 114°19'26.5"E) and Lamma Lo Tik Wan (LM, 22°13'12.9"N 114°07'39.5"E) are two of the 28 fish culture zones (HKAFCD, 2024a) governed by the Hong Kong Agriculture, Fisheries and Conservation Department (HKAFCD). Most of the farms are small family-run fish rafts, which collectively produce fewer than 500 tonnes of live fish – accounting for only 2% of local demand (HKAFCD, 2024a). Although mariculture is not a major source of seafood, the chemical, nutrient, and bacterial pollution produced by this industry impose considerable stress on the surrounding ecosystems (Lai et al., 2016). Both SK and LM are well sheltered by neighbouring bays and islands. LM on the west is more susceptible to inconsistent water quality influenced by the Pearl River while SK on the east is supplied by marine water from the South China Sea.

Sewage:

Center Island (CI, 22°26'14.1"N 114°13'16.6"E) is in the heart of Tolo Harbor, a slim coastal inlet of about 52 $km^2$ in area with a narrow bottleneck (~1 km). As a semi-enclosed embayment, water circulation is dictated by tidal movement which makes the area naturally prone to eutrophication (Chau, 2007). From the 80s to the late 90s, secondary treated sewage from an expanding population over 500 thousand was discharged into the harbor. In 1988 alone, there were over 40 episodes of harmful algal blooms recorded in the cove (HKEPD, 2007). Although the government had redirected the sewage discharge away since the late 90s, this area continues to record high levels of ammonia nitrogen, Escherichia coli, orthophosphate phosphorus, and biochemical oxygen demand compared to neighboring water bodies (HKEPD, 2023). Peng Chau (PC, 22°17'23.7"N 114°02'03.9"E) is better connected to the open sea by water currents than CI. However, Peng Chau Sewage Treatment Works, a secondary wastewater treatment facility with a design capacity of 3,250 $m^3$/day, discharges into this area via submarine outfall (HKEPD, 2001).

1.2 ARMS processing and molecular workflow

A typical set of data retrieved from an ARMS includes plate photos, taxonomic inventories of motile organism (>2mm), and molecular data from three bulk fractions (motile 500µm, motile 106µm, sessile). we focused on the data from the bulk samples. There are two motile fractions coming from sieving which followed the standard protocols (NMNH, n.d.-b, Figure S1). In brief, ARMS and ~60 L of surrounding seawater were collected and transported to a disassembly station in a bin. After all ARMS plates were disassembled, we filtered all the seawater together with the sediment and the motile organisms from the disassembly bin with a sieve set of 2 mm/500 µm and then a set of 500 µm/106 µm, which separated the motile organisms into two fractions: one between 2 mm and 500 µm (the motile 500 µm fraction), the other between 500 µm and 106 µm (the motile 106 µm fraction). About 15 ml of each motile fractions were rinsed with 90% ethanol before storage in a -20 °C freezer in 95% ethanol. The one sessile fraction came from organic materials that were collected from plate scraping following a different protocol (NMNH, n.d.-b): all sessile organisms were removed from the ARMS plate using a metal scraper, transferred into a blender, and homogenized. Similar to the motile fraction, ~ 15 ml sessile fraction was then preserved in 95% ethanol before storing in a -20 °C freezer.

DNA extraction and the downstream molecular workflow were identical for all three fractions except for an initial decanting step for the two motile fractions to remove sediments from the targeted organic matter. After this step, ~10 g material from each sample was weighed out and extracted from using the DNeasy PowerMax Soil Kit (Qiagen) using a slightly modified protocol (NMNH, n.d.-c) with an overnight chemical lysis with proteinase K rather than the standard physical lysis with glass beads aiming to enhance integrity of the DNA fragments. All DNA extracts were then purified using the DNeasy PowerClean Pro Cleanup Kit (Qiagen) with the standard protocol (QIAGEN, 2024).

We then targeted a 313 bp mitochondrial Cytochrome c oxidase I (COI/CO1) region using the "mlCOIintF/jgHCO2198" primer set for PCR amplification (Leray et al., 2013). All DNA samples were amplified in triplicate to minimize PCR bias. Each PCR reaction mixture contained 2 µl 10X Advantage 2 PCR Buffer (Takara Bio, USA), 1 µl of "mlCOIintF" and 1 µl of "jgHCO2198" (10 µM), 1.4 µl of 50X dNTP Mix (10 mM ea.), 0.4 µl 50X Advantage 2 Polymerase Mix, and 14.2 µl of a mixture of DNA and PCR grade of water (20 µl in total). Owing to natural variation of DNA quality, the final DNA input ranged from 10 ng ~50 ng for all samples. All samples were put through the same two-steps denaturing at 95 °C, followed by 16 cycles of 10 s denaturing at 95 °C -> 30 s annealing using a thermal cycling profile that started with an initial 1 min 62 °C -> 60 s elongation at 72 °C, then followed by 20 cycles of 10 s denaturing at 95 °C -> 30 s annealing at 46 °C -> 60 s elongation at 72 °C, then an extending elongation of 7 min at 72 °C, ending with a storage temperature of 4 °C. Triplicate amplifications were pooled as one sample before sequencing by the Genomics Core Centre for PanorOmics Science (CPOS) of the University of Hong Kong.

Two separate sequencing runs were conducted on the illumina MiSeq System using the MiSeq Reagent Kit v2 with 250-bp pair-end readings aiming for a minimum 100k pair-end reads per sample. All motile samples of the baseline/resistance phase ARMS were sequenced in one round

(32 samples in total), and the remaining samples were sequenced in another round (52 samples in total). All 84 samples (2 motile and 1 sessile fraction per ARMS, 28 ARMS total) were successfully sequenced.

#### 1.3 Bioinformatic pipeline

Sequences were first imported into QIIME2 (Bolyen et al., 2019) followed by adaptor removal using cutadapt (Martin, 2011) to clean the adaptor sequence used in PCR and NGS sequencing. Subsequently, DADA2 (Callahan et al., 2016) was used to first truncate sequences to 220 bp to filter out low quality reads, and then pair the forward and reverse reads, followed by denoising – a process removing potential errors, artifacts, and chimeras to construct amplicon sequence variants (ASVs). We removed ASVs detected in negative controls unless their sample abundance exceeded control levels by ≥ 10-fold. Since the targeted region (COI/CO1) is a protein coding region, we further filtered sequences based on amino acid translation. More specifically, MASCE (Ranwez, 2011) was used to first build an amino acid alignment based on the Moorea BIOCODE library. Then the invertebrate protein translation code was used to translate the remaining sequences to amino acid sequences. Afterwards, all translated sequences were aligned to the Moorea BIOCODE alignment removing any sequences that have stop codons, insertions, and more than three deletions (Leray et al., 2013). Lastly, VSEARCH (Rognes et al., 2016) was used to cluster the filtered ASVs into operational taxonomic units (OTUs) based on 97% sequence similarity.

Taxonomic assignment was conducted using BLAST within QIIME2 to a locally curated database (McIlroy et al., 2024). A minimum of 80% over at least 300 bp was considered a match to the phylum level. For all the unassigned OTUs, we performed a second round of BLAST against MIDORI (Leray et al., 2022), a regional COI database following the same criteria. Two assignment results were then combined for downstream analysis.

### 2. Supplementary Figures

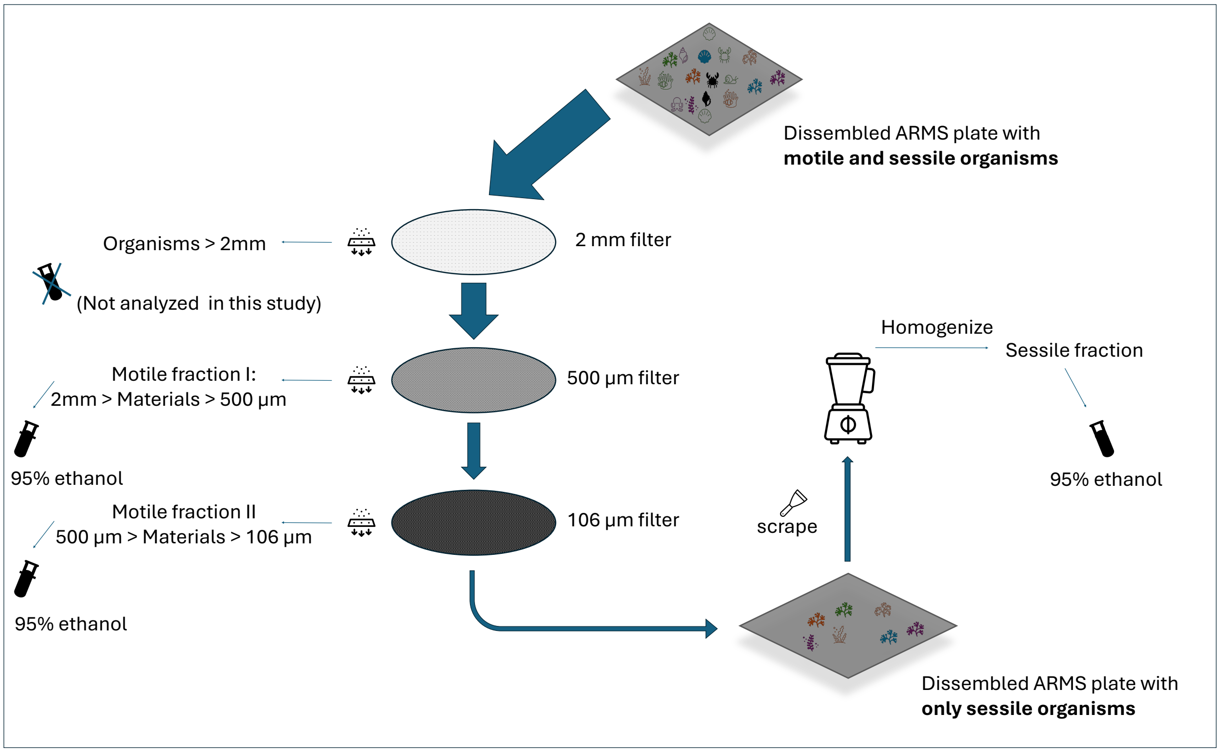


Figure S1. Post-dissembling, individual ARMS plates (top) were rinsed with filtered sea water to remove any motile organism and remaining debris. Dislodged materials were then sieved through a filter set of three (2 mm, 500 µm, 106 µm). Organisms over 2mm were not analyzed in this work. Motile fraction I and II were then preserved in 90% ethanol for later downstream processing. Sessile organisms retained on the plate were then scraped off and put though a blender to homogenize (sessile fraction), and then preserved with 95% ethanol.

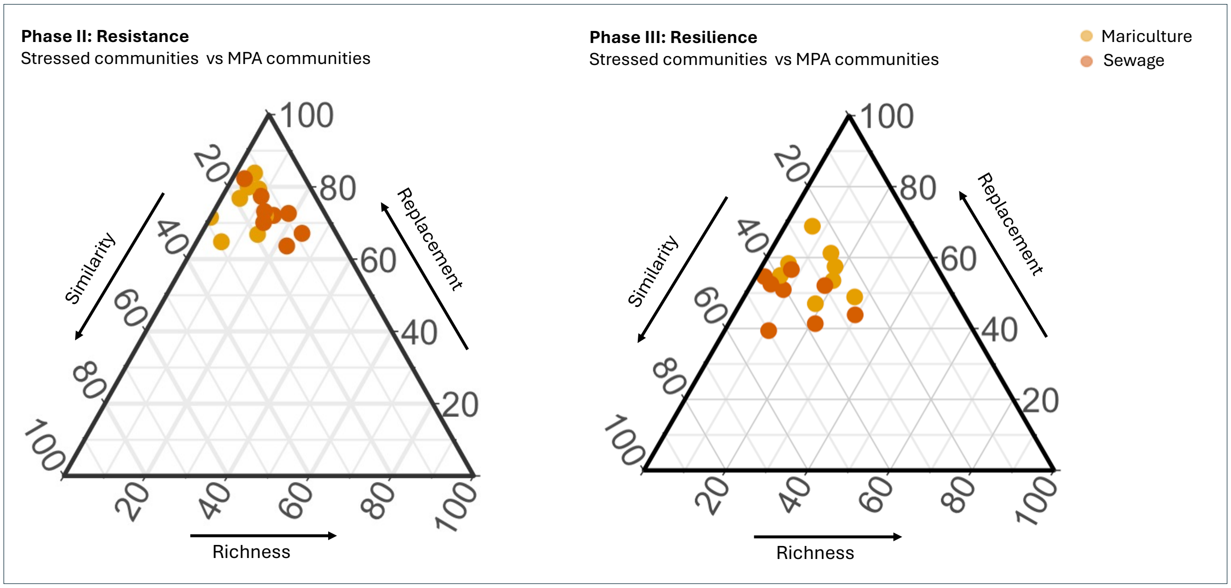


Figure S2. Ternary plots of beta diversity components (species replacement [βrp], richness difference [βrich]) during resistance (left) and resilience right) between stressed communities and MPA communities.

### 3. Supplementary Tables

Table S1. Treatment sites, their corresponding water monitoring stations and the GPS coordinates.

**Treatment Sites and Corresponding Water Monitoring Stations**

| | Site | Treatment | Latitude | Longitude | Monitoring Station | Latitude | Longitude |
|---|---|---|---|---|---|---|---|
| West | CDA | MPA | 22.20683 | 114.25672 | SM1 | 22.21230 | 114.23140 |
| | CDA | MPA | 22.20683 | 114.25672 | MM8 | 22.20035 | 114.32240 |
| | LM | Mariculture | 22.22025 | 114.12764 | SM3 | 22.22545 | 114.14970 |
| | LM | Mariculture | 22.22025 | 114.12764 | SM4 | 22.21263 | 114.13860 |
| | PC | Sewage | 22.28992 | 114.03442 | SM9 | 22.27367 | 114.06707 |
| | PC | Sewage | 22.28992 | 114.03442 | SM10 | 22.30208 | 114.03198 |
| | PC | Sewage | 22.28992 | 114.03442 | SM11 | 22.25738 | 114.01797 |
| East | TPC | MPA | 22.54292 | 114.43717 | MM5 | 22.52055 | 114.39388 |
| | SK | Mariculture | 22.36983 | 114.32403 | PM4 | 22.38233 | 114.31365 |
| | CI | Sewage | 22.43725 | 114.22183 | TM4 | 22.43273 | 114.21960 |

Table S2. Total species richness (top) and unique species (bottom) by fractions and by ARMS.

**Total OTUs and Unique OTUs by fractions and by ARMS**

| | | Phase I: Seeding, 12 months | | Phase II: Resistance, 24 months | | | | | | Phase III: Resilience, 30 months | | | | | |
|---|---|---|---|---|---|---|---|---|---|---|---|---|---|---|---|
| **Total OTUs** | | MPA: CDA | | MPA: CDA | | Mariculture: LM | | Sewage: PC | | MPA: CDA | | Mariculture: LM | | Sewage: PC | |
| West | ARMS No. | **89** | **90** | **79** | **81** | **83** | **57** | **88** | **61** | **77** | **78** | **84** | **58** | **87** | **62** |
| | 106 µm | 659 | 814 | 775 | 656 | 756 | 765 | 545 | 554 | 436 | 535 | 575 | 640 | 759 | 498 |
| | 500 µm | 153 | 256 | 385 | 234 | 262 | 252 | 222 | 362 | 216 | 170 | 184 | 253 | 200 | 492 |
| | Sessile | 626 | 565 | 529 | 604 | 544 | 386 | 426 | 408 | 356 | 466 | 626 | 683 | 515 | 488 |
| | By ARMS | 1209 | 1340 | 1249 | 1136 | 1136 | 1046 | 826 | 893 | 704 | 857 | 1078 | 1200 | 1137 | 1027 |
| | | MPA: TPC | | MPA: TPC | | Mariculture: SK | | Sewage: CI | | MPA: TPC | | Mariculture: SK | | Sewage: CI | |
| East | ARMS No. | **55** | **56** | **69** | **70** | **65** | **75** | **80** | **67** | **71** | **72** | **66** | **76** | **82** | **68** |
| | 106 µm | 771 | 799 | 747 | 829 | 855 | 967 | 605 | 735 | 581 | 592 | 754 | 665 | 603 | 686 |
| | 500 µm | 329 | 436 | 349 | 376 | 565 | 613 | 482 | 454 | 385 | 272 | 402 | 166 | 237 | 287 |
| | Sessile | 572 | 611 | 696 | 295 | 281 | 439 | 349 | 386 | 541 | 349 | 494 | 384 | 480 | 273 |
| | By ARMS | 1327 | 1432 | 1309 | 1124 | 1215 | 1428 | 968 | 1073 | 1101 | 914 | 1204 | 948 | 974 | 943 |

| | | Phase I: Seeding, 12 months | | Phase II: Resistance, 24 months | | | | | | Phase III: Resilience, 30 months | | | | | |
|---|---|---|---|---|---|---|---|---|---|---|---|---|---|---|---|
| **Unique OTUs** | | MPA: CDA | | MPA: CDA | | Mariculture: LM | | Sewage: PC | | MPA: CDA | | Mariculture: LM | | Sewage: PC | |
| West | ARMS No. | **89** | **90** | **79** | **81** | **83** | **57** | **88** | **61** | **77** | **78** | **84** | **58** | **87** | **62** |
| | 106 µm | 93 | 52 | 45 | 81 | 71 | 765 | 58 | 50 | 17 | 24 | 48 | 50 | 96 | 39 |
| | 500 µm | 9 | 14 | 40 | 17 | 3 | 252 | 8 | 30 | 10 | 5 | 12 | 16 | 21 | 24 |
| | Sessile | 26 | 71 | 36 | 68 | 34 | 386 | 28 | 18 | 21 | 37 | 49 | 59 | 48 | 33 |
| | By ARMS | 129 | 142 | 123 | 171 | 112 | 111 | 100 | 105 | 50 | 67 | 110 | 126 | 176 | 99 |
| | | MPA: TPC | | MPA: TPC | | Mariculture: SK | | Sewage: CI | | MPA: TPC | | Mariculture: SK | | Sewage: CI | |
| East | ARMS No. | **55** | **56** | **69** | **70** | **65** | **75** | **80** | **67** | **71** | **72** | **66** | **76** | **82** | **68** |
| | 106 µm | 49 | 53 | 96 | 95 | 103 | 116 | 57 | 89 | 27 | 30 | 63 | 55 | 34 | 58 |
| | 500 µm | 12 | 25 | 20 | 19 | 40 | 62 | 64 | 41 | 18 | 13 | 34 | 5 | 7 | 8 |
| | Sessile | 64 | 83 | 79 | 14 | 9 | 29 | 16 | 23 | 41 | 23 | 24 | 25 | 47 | 14 |
| | By ARMS | 125 | 161 | 198 | 136 | 156 | 217 | 145 | 162 | 89 | 68 | 127 | 88 | 89 | 81 |

Table S3. Z-scores and adjusted *p* values from Negative Binomial model to study community succession. Bold value highlighted in adjusted *p* indicated significant trends ($p < 0.05$ after Benjamini-Hochberg adjustment).

### Z score and adjusted *p* - value from Negative Binomial model to assess succession

| phylum | MPA | slope | z score | Adj *P* value |
|---|---|---|---|---|
| Arthropoda | CDA | -3.04E-02 | -3.92 | **4.05E-04** |
| Annelida | CDA | -2.86E-02 | -3.65 | **7.46E-04** |
| Mollusca | CDA | -2.29E-02 | -1.49 | 1.46E-01 |
| Bacillariophyta | CDA | -3.94E-02 | -1.59 | 1.30E-01 |
| Rhodophyta | CDA | -2.77E-02 | -3.13 | **3.27E-03** |
| Porifera | CDA | -2.69E-02 | -3.12 | **3.27E-03** |
| Total richness | CDA | -2.37E-02 | -3.11 | **3.27E-03** |
| Arthropoda | TPC | -3.44E-02 | -7.83 | **6.64E-14** |
| Annelida | TPC | -2.71E-02 | -5.19 | **1.47E-06** |
| Mollusca | TPC | -3.16E-02 | -2.94 | **4.72E-03** |
| Bacillariophyta | TPC | 2.79E-02 | 2.93 | **4.72E-03** |
| Rhodophyta | TPC | 1.12E-02 | 1.79 | 9.40E-02 |
| Porifera | TPC | 8.39E-03 | 1.05 | 2.92E-01 |
| Total richness | TPC | -1.64E-02 | -3.68 | **7.46E-04** |

Table S4. Mean values of all 13 environmental parameters over the study period (Jun 20 ~ Dec 22) from two marine park (CDA, TPC), and adjusted *p*, adjusted $R^2$ value from linear model (all df = 1,131). Parameters showed significant site differences were highlighted in blue ($p < 0.05$ after Benjamini-Hochberg adjustment).

**30 months' mean environmental parameters of two MPA, and the Adj *p* value and Adj R2**

| | CDA | TPC | Adj *P* value | AdjR2 |
|---|---|---|---|---|
| Nitrite Nitrogen (mg/l) | 0.012 | 0.004 | **3.51E-03** | 8.98% |
| Total Inorganic Nitrogen (mg/l) | 0.092 | 0.048 | **5.07E-03** | 7.59% |
| Nitrate Nitrogen (mg/l) | 0.055 | 0.018 | **8.48E-03** | 6.37% |
| Orthophosphate Phosphorus (mg/l) | 0.007 | 0.005 | **3.97E-02** | 3.97% |
| Phaeo pigments (μg/L) | 0.75 | 0.36 | 5.86E-02 | 3.18% |
| Suspended Solids (mg/L) | 4.08 | 2.90 | 1.32E-01 | 1.91% |
| Chlorophyll a (μg/L) | 3.16 | 1.77 | 2.98E-01 | 0.75% |
| Dissolved Oxygen (mg/L) | 5.92 | 5.76 | 5.77E-01 | -0.25% |
| Turbidity (NTU) | 5.91 | 7.39 | 5.77E-01 | -0.13% |
| *E. coli* (cfu/100ml) | 1.38 | 0.70 | 5.77E-01 | -0.31% |
| Dissolved Oxygen saturation (% of saturation) | 84.89 | 83.46 | 6.83E-01 | -0.52% |
| Faecal Coliforms (cfu/100ml) | 2.99 | 1.91 | 7.34E-01 | -0.63% |
| Ammonia Nitrogen (mg/l) | 0.023 | 0.024 | 7.60E-01 | -0.69% |
